# FvLcp1, a type-D fungal LysM protein with Chitin-binding domains, is a secreted protein involved in host recognition and fumonisin production in *Fusarium verticillioides* - maize kernel interaction

**DOI:** 10.1101/789925

**Authors:** Huan Zhang, Man S. Kim, Jun Huang, Huijuan Yan, Tao Yang, Linlin Song, Wenying Yu, Won Bo Shim

## Abstract

- *Fusarium verticillioides* is one of the key maize ear rot pathogens and produces fumonisins, a group of mycotoxins detrimental to humans and animals. Unfortunately, our understanding on how this fungus interacts with maize kernels to trigger mycotoxin biosynthesis is very limited.
- We performed a systematic computational network-based analysis of large-scale *F. verticillioides* RNA-seq datasets to identify potential gene subnetwork modules that are associated with virulence and fumonisin regulation.
- Among the highly discriminative subnetwork modules, we identified a putative hub gene *FvLCP1*, which encodes a putative a type-D fungal LysM protein with a signal peptide, three LysM domains, and two chitin binding domains. FvLcp1 is a unique protein that harbors these domains amongst five representative *Fusarium* species.
- FvLcp1 is a secreted protein important for fumonisin production with LysM domain playing acritical role. Chitin-binding domain was essential for *in vitro* chitin binding.
- Using rice blast fungus, we learned that FvLcp1 accumulates in appressoria, a key infection structure, suggesting that FvLcp1 could be involved in host recognition and infection. Also, full length FvLcp1 was able to suppress the BAX triggered plant cell death in *Nicotiana benthamiana*.
- This is the first report where a unique type-D LysM secreted protein with chitin-binding domain in mycotoxigenic fungus *F. verticillioides* was shown to be potentially involved in suppressing host cell death and promoting fumonisins biosynthesis while the pathogen colonizes maize kernels.

## INTRODUCTION

*Fusarium verticillioides* (telemorph *Gibberella moniliformis*) is a prevalent fungal pathogen associated with maize wherever the crop is grown. Not only does the pathogen cause yield-limiting maize ear rot and stalk rot diseases, it also produces a group of carcinogenic mycotoxins fumonisins on infested ears that pose serious health risk to animals and humans (Gelderblom *et al*., 1992; Bacon & Nelson, 1994; Munkvold & Desjardins, 1997; Marasas, 2001). This family of structurally-related mycotoxins contaminate maize and maize-based products worldwide (Chulze *et al*., 1998; Castella *et al*., 1999; Gutema *et al*., 2000; Logrieco *et al*., 2002; Arino *et al*., 2007), and fumonisin B1 (FB_1_) is the most commonly form found in nature with highest toxicity. In addition to potential health risks to humans, fumonisin consumed by animals causes diseases such as leukoencephalomalacia in horses and pulmonary edema in pigs (Thiel *et al*., 1991; Osweiler *et al*., 1992; Badria *et al*., 1996; Gelderblom *et al*., 2001; Voss *et al*., 2001). Economic loss and food safety risks associated with maize contaminated with fumonisins have been documented in numerous publications (Wu, 2004; Wang *et al*., 2006; Wu, 2006; Wu *et al*., 2011). There are several factors that make it extremely difficult to predict potential outbreak of fumonisins in maize, mainly due to different cultural practices, abiotic epidemiological factors, and pathogen biology. Durable maize resistance against *F. verticillioides* and fumonisin contamination is not presently available to growers, and current fumonisin mitigation tools are few and only partially effective at best. Fumonisin contamination occurs during late stages of maize ear development, making it difficult for growers to apply intervention strategies.

There is a critical need to better understand the mechanism of fumonisin biosynthesis and regulation in *F. verticillioides* to develop effective long-term control strategies. Recent studies have documented the intricate transcriptional and epigenetic regulation that affects mycotoxin gene clusters and how these molecular networks that respond to environmental factors are linked to genetic components regulating mycotoxin production (O’Brian *et al*., 2007; Georgianna & Payne, 2009; Woloshuk & Shim, 2013). Whole genome sequences provide reference database for genomic and transcriptomic analyses, and these efforts have improved our understanding of tissue-specific colonization and fumonisin production (Shim *et al*., 2003; Bluhm & Woloshuk, 2005; Ma *et al*., 2010; Shu *et al*., 2011). Furthermore, recent technological advancements in next generation sequencing (NGS) and computational biology are facilitating new discoveries in gene regulatory networks (Yoon & Qian, 2009; Sahraeian & Yoon, 2011). In our earlier studies (Kim *et al*., 2018a; Kim *et al*., 2018b), we demonstrated how network-based comparative analysis of transcriptome data through probabilistic subnetwork inference can help identify potential pathogenicity-associated subnetwork modules in *F. verticillioides*. Plant-microbe associations come in many different schemes, due to the intricate evolutionary relationship between the host and the pathogen (Jones & Dangl, 2006; Spoel & Dong, 2012; Dangl *et al*., 2013). Pre-harvest rots occurring in crops are predominantly caused by fungal organisms, namely by pathogens having the ability to decompose and putrefy plant tissues. In a similar context, it is reasonable to hypothesize that ear rot fungal pathogens have also developed specialized mechanisms for host adaptation, virulence, and mycotoxin biosynthesis.

One intriguing question we asked in this study was whether secreted proteins play a role in triggering fumonisin production in *F. verticillioides* when the fungus interacts with maize kernels. In recent years, studies on how fungal secreted proteins play critical roles in triggering host defense or susceptibility are innovating our understanding of various plant-fungal interactions (de Jonge *et al*., 2010; Dangl *et al*., 2013; Liu *et al*., 2013; Dong *et al*., 2015; Sharpee *et al*., 2017). Thus, we modified our computational network analysis strategy to primarily select genes that encode secreted proteins to construct our subnetwork modules. Through this approach, we identified *FvLCP1* that was predicted as a hub gene in a robust subnetwork. *FvLCP1* encodes a putative secreted protein with two functional domains, LsyM and chitin-binding domains, that are known to trigger host responses. We further characterized the role of FvLcp1 in FB_1_ production through motif deletion experiments, followed by protein secretion, chitin-binding activity and subcellular localization assays to gain a deeper understanding of FvLcp1. Lastly, we used *Magnaporthe oryzae -* rice and *Agrobacterium tumefaciens* - *Nicotiana benthamiana* systems to test whether FvLcp1 is associated with host recognition and suppression of host cell death, respectively.

## MATERIALS AND METHODS

### RNA sample preparation, sequencing and data preprocessing

The wild-type strain *Fusarium verticillioides* 7600 (Ma *et al*., 2010) was cultured at 25°C on V8 juice agar (200 ml of V8 juice, 3 g of CaCO3 and 20 g of agar powder per liter). Surface sterilized kernels of maize inbred line B73 (provided by USDA-ARS, Iowa State University) and hybrid 33K44 (provided by Pioneer Hybrid) were inoculated with *F. verticillioides* as described previously (Ortiz & Shim, 2013; Shin *et al*., 2013) and samples were collected at 2, 4, 6, and 8 days post inoculation (dpi). For each replicate, 10 inoculated kernels were pooled, and total RNA from kernels was extracted with kernel extraction buffer (50Mm TRIS pH 8.0, 150mM LiCl, 5mM EDTA pH8.0, 1% SDS) for high polysaccharide rich tissue. Kernel extraction buffer (200 µl) was combined with 200 µg of ground-frozen seed tissue and 200 µl phenol: chloroform (pH 4.3). The mixture as vortexed until thawed and incubated on ice for 5 min. The mixture was then transferred into PHASELOCK tubes (5 PRIME, Hamburg, Germany) for phase-separation based purification and centrifuged. Phenol: chloroform (200 µl, pH 4.3) was added, vortexed, and mixture was incubated on ice for 5min, and centrifuged. Afterwards, RNA was purified with RNeasy Mini Spin Column in RNeasy Mini Kit (Qiagen, Hilden, Germany) according to manufacturer’s specifications. Then RNA was DNAase treated for sequencing. RNA samples from uninoculated B73 and 33K44 kernels (0 day) were also collected and prepared as negative controls. We collected eight pooled samples for each time point. The RNA samples were preprocessed for QA/QC, and library construction was performed following the standard protocol at Texas A&M AgriLife Genomics and Bioinformatics Services (College Station, Texas). Using Illumina HiSeq 2500, paired-end RNA sequencing was conducted, thereby generating 125-bp reads for *F. verticillioides* inoculated on two different maize kernels (*i.e.* hybrid 33k44 vs. inbred B73). Through this sequencing process, fourteen independent sequenced RNA libraries (*i.e.*, seven biological replicates and seven technical replicates) at two time points (6 dpi and 8 dpi) were generated for the two maize kernel samples. Therefore, 56 paired-end RNA libraries were prepared in total.

We first preformed preprocessing the RNA-seq datasets by alignment using TopHat2 (Kim *et al*., 2013) along with HTSeq (Anders *et al*., 2015), filtering out relatively insignificantly expressed genes, and performing a two-step normalization (Kim *et al*., 2018a). Therefore, three different preprocessed gene expression matrices (PGEM); i) PGEM-1: RPKM-control; ii) PGEM-2: RPKM-TMM; iii) PGEM-3: TPM-TMM were obtained (Mortazavi *et al*., 2008; Robinson *et al*., 2010; Wagner *et al*., 2012). Beta-tubulin genes were utilized as control genes.

To search for genes encoding secreted protein, we assigned genes with signal peptide that were significantly differentially expressed between the two maize kernels as seed genes by measuring *t*-test statistics scores as well as F scores of ANOVA across all three PGEMs, thereby preparing ten seed genes for each kernel (twenty in total). For co-expression network construction, we built five different networks at five different levels (*i.e.*, five different threshold cut-offs) as previously applied (Kim *et al*., 2015; Kim *et al*., 2018a; Kim *et al*., 2018b), where the smallest size included roughly 400,000 edges and the largest size contained around 2,000,000 edges. Additional detail is provided in Supplementary Method A.

### Subnetwork modules extension

By following our previously proposed analysis approach (Kim *et al*., 2018a), we started extending subnetwork modules having member genes up to two from the seed genes (Fig. 1). For instance, from a seed gene on *F. verticillioides* co-expression networks based on a PGEM, we extend up to three subnetwork modules as long as i) their discriminative power increase (measured by *t*-test statistics) between the two maize kernels (B73 vs 33K44) exceeds 5% and ii) the difference in discriminative power increase between two suboptimal modules and the optimal module with the highest discriminative power is less than 2%. When searching for the third member gene, we started applying computationally efficient adaptive branch-out technique while adjusting stopping criterion (MDPE: Minimum Discriminative Power Enhance) to find candidate subnetwork modules as proposed in this approach. As shown in Fig. 1, we start adaptive branch-out with the desired stopping criterion at first by 10%, and continuously branching out modules when the following requirements of a candidate subnetwork module are satisfied; a) relatively high discriminative power; b) relatively large number of edges of the seed gene; c) significant GO term annotation; and d) distance between the seed gene and all member genes less than five.

**Figure 1.**
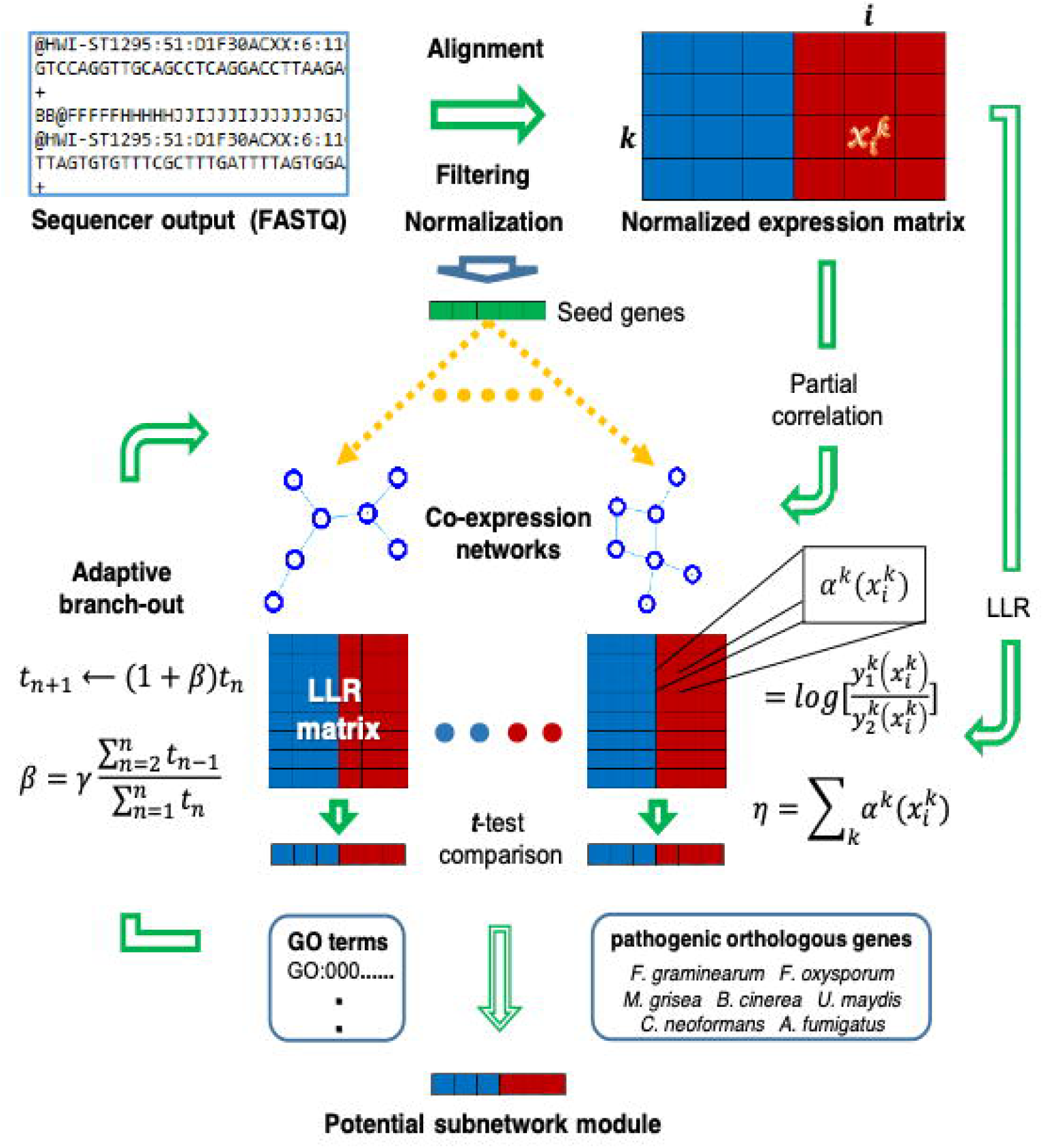
Schematic overview of the proposed network-based comparative analysis approach to identify potential subnetwork modules associated with the *F. verticillioides* pathogenicity. The normalized gene expression matrix is obtained from the sequencer output through preprocessing such as mapping, filtering, and normalization. Based on the preprocessed gene expression matrix, we constructed six co-expression networks threshold at six different partial correlation levels and also prepared a log-likelihood ratio (LLR) matrix. Next, we extended subnetwork modules from seed genes, significantly differentially expressed genes between the two conditions, utilizing a computationally efficient adaptive branch out approach. In order to identify robust potential subnetwork modules, we adaptively adjusted parameters when branching out the subnetwork modules by considering whether at least one known orthologous gene in other fungi is impactful in a module and also the member genes of the module are annotated by significant GO terms. Finally, each potential subnetwork module was selected if the module demonstrates not only strong association with the pathogenicity of the fungi, but also significantly differentiated activity levels between the two conditions.

If a subnetwork module meets requirements a), b), and c) but disagrees with d), we then start raising the desired stopping criterion by 1% at a time and continuously perform the same approach as long as either all the requirements are not satisfied yet or a subnetwork module failed to reach six members (minimum number of module member genes). For every seed gene in all co-expression networks based on all three PGEMs, we continuously repeated the adaptive branch-out process to identify candidate functional subnetwork modules.

After collecting all possible subnetwork modules associated with signaling activity, we listed seed genes whose subnetwork modules were vigorously activated across all three PGEMs. Among the list of genes, we selected a representative module with the highest *t*-test statistics score at each PGEM (three modules in total), and subsequently performed module combining across PGEMs and post-pruning process. In order to only keep genes relatively further associated with the member genes including particularly their seed genes, we removed genes if they don’t meet the following conditions; i) capability to help their modules differentiate relatively more between the two strains; ii) strong association with other genes including their seed genes across different PGEMs; iii) functional relevance with other member genes (*i.e.*, GO terms). Additional detail provided in Supplementary Method B∼F.

### Nucleic acid manipulation, polymerase chain reaction (PCR), and transformation

Fungal strains were grown in YEPD liquid medium (Difco, Detroit, MI), and genomic DNA was extracted using an OmniPrep Genomic DNA Extraction kit (G Biosciences, St. Louis, MO). The constructs for transformation were generated by split-marker approach via homologous recombination and transformed into wild-type protoplast as described previously (Sagaram & Shim, 2007). To further investigate the functional role of different FvLcp1 domains, we first generated four complementation constructs via homologous recombination; the first one has a complete deletion of signal peptide (ΔSP), the second one has a complete deletion of LysM domains (ΔLysM), the third one has a complete deletion of ChtBD1 domains (ΔChtBD1), and the fourth one is complete wild-type genes (FvLCP1C). These constructs were all driven by its native promoter and terminator. Additional detail provided in Supplementary Method G. Transformants were regenerated and selected on regeneration medium containing 100 µg/ml of hygromycin B (Calbiochem, La Jolla, CA, USA) and/or 150 µg/ml G418 sulfate (Cellgro, Manassas, VA, USA) as needed. Respective drug-resistant colonies were screened by PCR and qPCR for the presence and expression of transformed constructs, respectively (data not shown). All primers used in this study are listed in Supplementary Method H.

For qPCR assay, total RNA samples were prepared by using RNeasy plant mini kit (Qiagen) according to manufacturer’s specifications and quantified using Nanodrop.

Subsequently, total RNA samples were converted into cDNA using the Verso cDNA synthesis kit (Thermo Fisher Scientific, Waltham, MA) following the manufacturer’s protocol. qRT-PCR analyses were performed using the SYBR Green Dynamo Color Flash qPCR kit (Thermo Fisher Scientific) on an Applied Biosystems 7500 Real-Time PCR system.

### Maize kernel infection, FB_1_ and ergosterol analyses

FB1 and ergosterol HPLC analyses were performed as previously described (Ortiz & Shim, 2013; Shin *et al*., 2013). FB_1_ and ergosterol quantifications were performed by comparing HPLC peak areas to FB_1_ and ergosterol standards, respectively (Sigma, St, Louis, MO, USA). FB_1_ levels were then normalized to ergosterol contents in samples by calculating [FB_1_ ppm / ergosterol ppm] x 100.

### Similarity analysis between LysM domains

Forty-nine LysM protein sequences from fungi and plants were obtained from NCBI database and verified by SMART (http://smart.embl-heidelberg.de/) (Letunic *et al*., 2011; Liu, B *et al*., 2012; Kombrink *et al*., 2017). Subsequently, we used ClustalX for sequence alignment, and similarities scores were calculated by SIAS (Sequence Identity And Similarity) (http://imed.med.ucm.es/Tools/sias.html). The calculated similarity scores were displayed in R, and LysM domain sequence logos were created by Skylign tool (http://skylign.org) (Wheeler *et al*., 2014).

### Yeast secretion trap assay

The YS-0 vector (Lee *et al*., 2006) which carries a truncated invertase gene *SUC2* that lacks the first two amino acids (Met21 and Thr22) of the mature protein was used. The full-length open reading frames (ORFs) with or without the signal peptides were cloned into pYST-0 vector which were previously had been digested with *EcoR*I or *Not*I. YST constructs were transformed into an invertase secretion-deficient yeast strain YTK12 (provided by Drs. Guo-Liang Wang and Wende Liu, Chinese Academy of Agricultural Science, Beijing, China), and plating the transformants on a medium containing sucrose in place of glucose as the sole carbon source at 30 °C. All transformants were restreaked on sucrose agar plates and confirmed by PCR with gene-specific primers.

### Generation of the FvLcp1-GFP fusion construct and microscopy

To generate the FvLcp1-GFP fusion construct, a 4,009-bp DNA fragment containing the native promoter and entire FvLCP1 gene sequence were amplified by primer pair LysM_GFPF/R. The 4,009-bp PCR product was inserted into the *Xho*I and *Cla*I sites of pKNTG vector which harbors a GFP tag and the geneticin-resistance marker (Zhong *et al*., 2015). The resulting construct was confirmed by sequencing and then transformed into *F. verticillioides* wild-type strain 7600 and *M. oryzae* wild-type strain Guy11. The presence of geneticin-resistance marker in transformants were confirmed by PCR.

To visualize the subcellular localization of FvLcp1 in the different vegetative growth stages of *F. verticillioides*, collected spores were incubated in YEPD medium for 12-24 hours at 28 ℃ at 180 rpm. Subsequently, conidia, germinated conidia, and vegetative hyphae were observed under the Nikon A1 laser scanning confocal microscope as previously described (Zheng, W *et al*., 2016). To observe the vacuolar membrane or vacuole, conidia, germinal conidia, and vegetative hyphae were treated with FM4-64 or CMAC as previously described (Zheng *et al*., 2015). The collected spores (5 × 10^4^ conidia/mL) of *M. oryzae* mutant Guy11/FvLcp1-GFP were inoculated into the rice sheath of susceptible rice cultivar CO39 as previously described (Zheng, H *et al*., 2016). After inoculation for 12 hours, 24 hours and 36 hours, samples were examined using an Olympus-BX51 fluorescence microscope.

### Chitin Binding assay

The sequences of mature FvLcp1 (without signal peptide) and truncated FvLcp1 (ΔLysM truncated and ΔChtBD1 truncated) were amplified from *F. verticillioides* wildtype 7600 cDNA and inserted into pGEX-KG which containing a GST-tag and an ampicillin-resistant selective marker by One-Step Cloning Kit (Vazyme, Nanjing, China) according to the manufacturer’s instructions. The fragments were then transferred into *E. coli* (DH5-alpha) and confirmed by PCR and sequencing. After sequencing verification, we transformed the plasmids into *E. coli* Rosetta competent cells (Novagen, Madison, WI) for protein expression. After confirmation by PCR, we cultured the selected colonies in LB medium at 37℃ until the value of an optical density at 600 nm (OD600) reached 0.4. The heterologous production of mature FvLcp1 and truncated FvLcp1 were induced with 0.1M IPTG at 28℃ for approximately 6 hours. Harvested bacterial cultures (100 mL) were lysed with 5-7 mL lysis buffer (50 mM NaCl, 20 mM Tris-HCl, 1 mM EDTA, 0.05% Triton X-100, 1 mM PMSF, and appropriate protease inhibitor tablets).

After lysis in 5 second on/off cycles for about 15 minutes in UP-250 portable ultrasonic cell grinder (Ningbo XinZhi, Ningbo, China), the lysates were centrifuged at 13000-14000 g for 30-45 minutes at 4℃. Chitin magnetic beads (New England Biolabs, Ipswich, MA) were used for investigating the affinity of mature FvLcp1 and truncated proteins. Proteins supernatants and chitin beads were incubated at 4℃ overnight with agitation in 5 ml of lysis solution. Following SDS-PAGE, the chitin-binding ability of mature FvLcp1 and truncated FvLcp1 proteins were examined in the insoluble pellet fraction by western blot using anti-GST (1:5000) as primary antibody and Goat anti-Mouse IgG (1:10000) as secondary antibody.

### Maize kernel and rice leaf virulence assay

Maize kernel infection assay was performed as described previously (Mukherjee *et al*., 2011). Briefly, maize kernels B73 and 33K44 were surface sterilized, and fungal spore droplets were applied to wound sites created with a syringe needle. Fungal colonization of seeds were monitored after one-week incubation. For rice infection, we followed the method described before (Zhong *et al*., 2015). Briefly, *M. oryzae* wild-type (Guy11) or Guy11/FvLcp1-GFP transformants were grown in rice bran medium (2% rice-polish, 1.5% agar, pH 6.0) for about 7-10 days until produced enough conidia. Diluting the concentration of conidia suspension to approximate 1×10^5^ spore/mL in 0.02% Tween 20 solution and sprayed on susceptible rice cultivar CO39.

### Gateway cloning and *Agrobacterium tumefaciens* infiltration assays

The mature sequence of FvLcp1 without its signal peptide was first amplified by PL4F/R then cloned into the donor vector, pDONR201, after confirmed by sequencing, then transferred into the destination vector pGWB505 which containing the 35S promoter by the gateway cloning using described methods (Nakagawa *et al*., 2007). The LysM domain deletion mutant was constructed by primer pairs PL4F/01584-LysM-R and 01584-LysM-F/PL4R, then fused together by single-joint PCR. Similarly, the ChtBD1 domain deletion mutant was generated by PF4F/01584-Chitin-R and 01584-Chitin-F/PF4R, then fused together by single-joint PCR. Following the method described above, LysM domain deletion construct and ChtBD1 domain deletion construct were cloned into pDONR201, then transferred into pGWB505 after sequencing. The resulting constructs were transformed into *Agrobacterium tumefaciens* strain GV3101 cells. Individual transformants were screened by 30 µg ml^-1^ of rifampicin and 100 µg ml^-1^ spectinomycin then confirmed by PCR. We infiltrated tobacco *N. benthamiana* leaves as previously described (Li *et al*., 2009; Pitino *et al*., 2016).

First, *A. tumefaciens* containing FvLcp1 related constructs (mature FvLcp1, FvLcp1ΔLysM and FvLcp1ΔChtBD1) were infiltrated in the leaves of *N. benthamiana*, respectively. To test their ability to suppress BAX-triggered plant cell death, *A. tumefaciens* with BAX were infiltrated in the previous infiltrated area 24 hours after the first infiltrations mentioned above. Infiltrated leaves of *N. benthamiana* were underlined with black cycles. RNA samples were extracted by Eastep® Super Total RNA Extraction Kit (Promega, Madison, WI) by following the suggested protocol within 24-36 hours after infiltration. We used Takara Primescript^TM^ reagent Kit with gDNA eraser for the cDNA synthesis. Primers used in this experiment are listed in Supplementary Method H.

## RESULTS

### RNA-Seq data and subnetwork module establishment

In order to systematically analyze transcriptional activity of functional *F. verticillioides* genes that encode proteins with signal peptide, we performed network-based comparative analysis with adaptive branch-out technique using three different preprocessed gene expression matrices (PGEMs). We obtained paired-end sequencing reads by capturing transcriptomes of *F. verticillioides*-inoculated maize inbred B73 and hybrids 33k44 kernels at two time points (6 dpi & 8 dpi). Through preprocessing, 14.7% of *F. verticillioides* were removed due to their insignificant expression levels, and therefore 13,533 genes were utilized for downstream analysis. Table S1 provides general information of both RNA-Seq datasets (*F. verticillioides* on 33K44 vs B73). Based on the three PGEMs, we inferred co-expression networks of *F. verticillioides* on both B73 and 33K44. Co-expression network sizes, from the smallest (400,000 edges) to the largest (2,000,000 edges), distributed by threshold cut-offs (partial correlation coefficients) are listed in Table S2 A and B. Since our main objective was to search for functional subnetwork modules that were differentially activated on co-expression networks across all three PGEMs, we also considered how co-expression networks from different PGEMs were similar to each other. Table S3 demonstrates degree of similarity between co-expression networks by calculating percentages of number of the same edges shown on two different networks from two different PGEMs.

We selected ten *F. verticillioides* genes that show significant differential expression in B73 or 33K44 as seed genes. We considered *t*-test scores and F scores of ANOVA, so that every seed gene had certain minimum threshold (*t-*test score >= 4 and F score >= 4) for at least one PGEM (Table S4). As we extended subnetworks modules from the seed genes by satisfying the our computational requirements (Supplementary Method D &E), we acquired three seed genes whose modules were significantly activated across all three PGEMs for each maize kernels (FVEG_01584, FVEG_07670, and FVEG_11722 in B73; FVEG_00397, FVEG_01726, and FVEG_05574 in 33K44) as shown in Table S5. From these six subnetwork modules, we further focused on top two modules from each strain after considering averaged *t*-test scores of representative modules of each PGEM. Therefore, representative modules based on FVEG_01584 & FVEG_07670 were selected to be potential subnetwork modules for B73 strain (Fig 2A & B). For 33K44 strain, modules of FVEG_00397 & FVEG_05574 were chosen to be representatives (Fig 2C & D). Representative modules of FVEG_01584 & FVEG_07670 showed higher *t*-test scores than that of FVEG_11722 by 19.5% and 6.0% while modules of FVEG_00397 & FVEG_05574 demonstrated higher *t*-test scores than that of FVEG_01726 by 7.0% and 5.8%.

**Figure 2.**
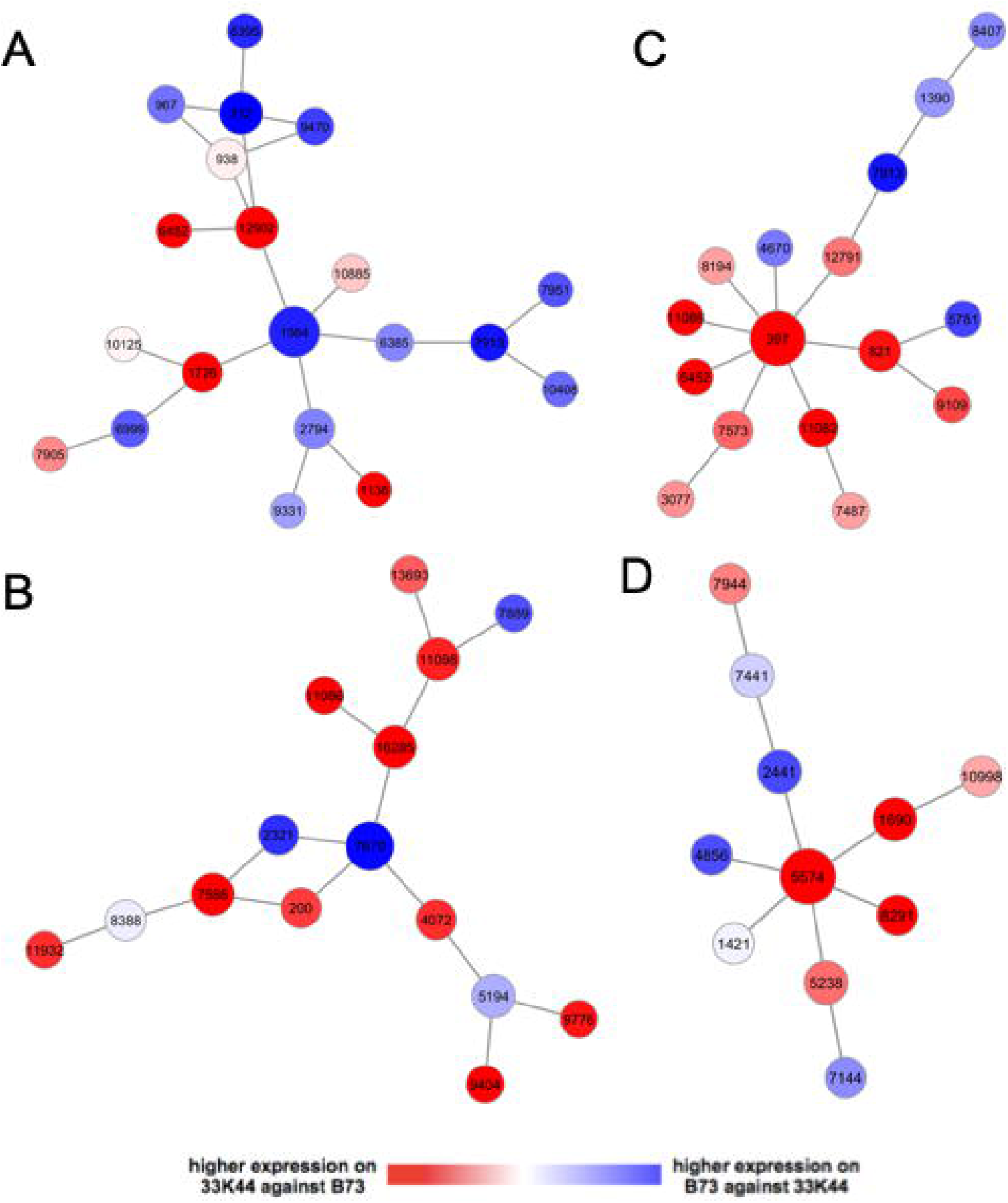
Potential *F. verticillioides* subnetwork modules associated with 33K44 against B73. (A) FVEG_01584 module: three subnetwork modules (size-2 from PGEM-1, size-4 from PGEM-2, and size-5 from PGEM-3) were combined and 31% of them were eliminated through the post-pruning process. (B) FVEG_07670 module: integration of three modules (size-4 from PGEM-1, size-2 from PGEM-2, and size-4 from PGEM-3) was carried out and 46% of them were removed. (C) FVEG_00397 module: three modules (size-5 from PGEM-1, size-5 from PGEM-2, and size-1 from PGEM-3) were merged and 33% of them were excluded; (D) FVEG_05574 module: incorporation of three modules (size-3 from PGEM-1, size-2 from PGEM-2, and size-2 from PGEM-3) completed and 27% of them were ruled out.

### FVEG_01584 gene (*FvLCP1*) encodes a protein with unique domain composition amongst Fusarium species

Among four subnetworks identified through our computational pipeline, we selected the subnetwork module with the predicted hub FVEG_01584 for further study. This subnetwork has sixteen member genes showing strong correlation, with a mixture of genes showing differential expression in B73 and 33K44 maize. Majority of the genes identified in the subnetwork encode uncharacterized or hypothetical proteins. The predicted hub gene FVEG_01584, which we designated *FvLCP1* (*<u>F</u>usarium <u>v</u>erticillioides* <u>L</u>ysM-<u>C</u>ontaining <u>P</u>rotein <u>1</u>), encodes a putative 733-amino-acid secreted protein containing an N-terminal signal peptide, three LysM domains and two chitin-binding (ChtBD1) domains (Fig. 3). We queried FvLcp1 protein sequence against the protein databases of *Fusarium verticillioides* 7600 (taxid:334819), *Fusarium graminearum* PH-1 (taxid:229533), *Fusarium oxysporum f. sp. lycopersici* 4287 (taxid:426428), *Fusarium fujikuroi* IMI 58289 (taxid:1279085) and *Nectria haematococca* mpVI 77-13-4 (taxid:660122) (Ma et al., 2010, Coleman et al., 2009, Wiemann et al., 2013), and learned that FvLcp1 is a unique protein that harbors both LysM and chitin binding domains in these five representative *Fusarium* species (Fig. 3). This outcome led us to question whether FvLcp1 plays a distinct role in *F. verticillioides*, particularly during maize infection and fumonisin biosynthesis.

**Figure 3.**
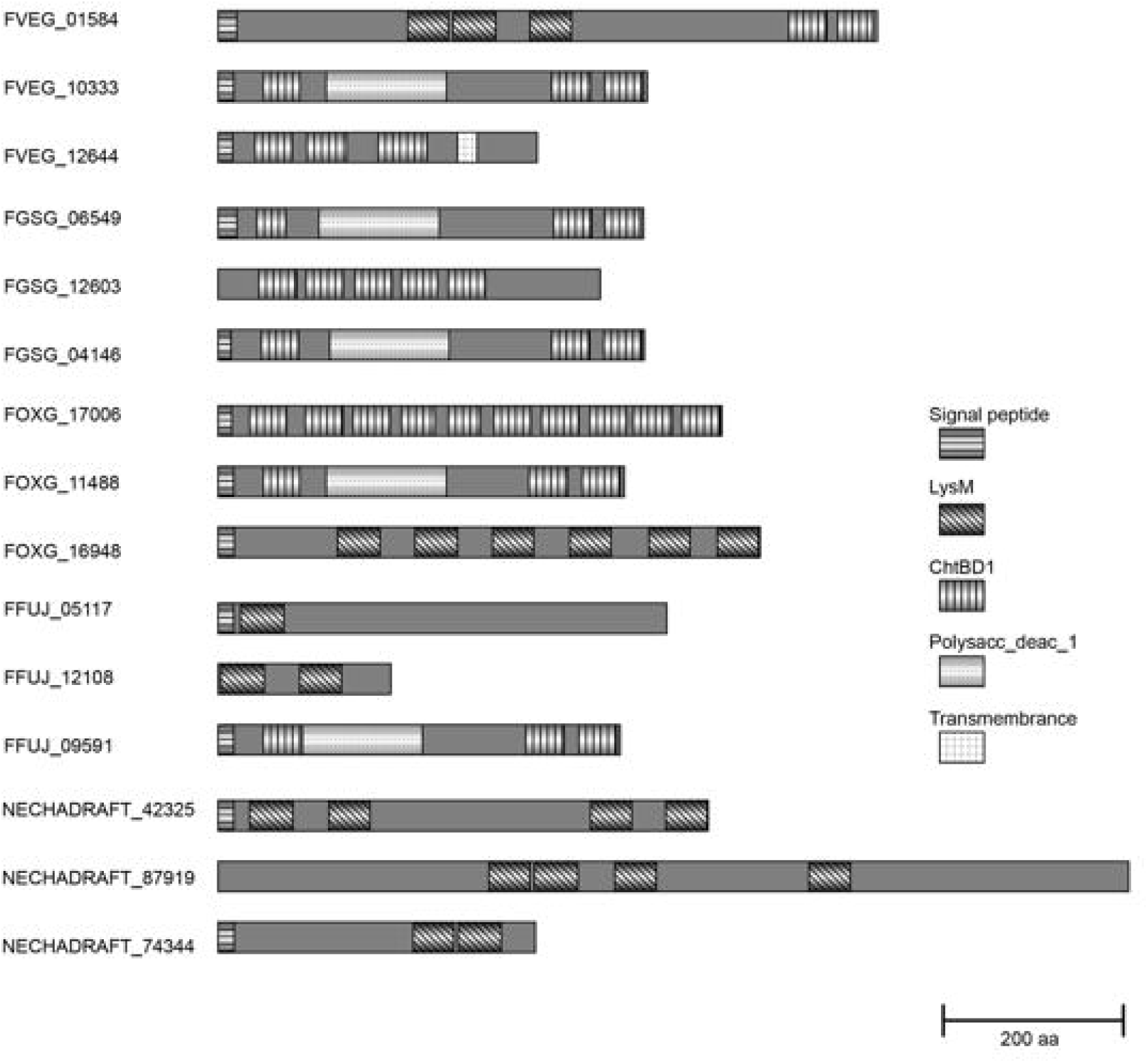
Domain analysis of FvLcp1 against other similar proteins identified in key plant pathogenic *Fusarium* species. Putative paralogs and orthologs of FvLcp1 in *F. verticillioides*, *F. graminearum*, *F. oxysporum* f.sp. *lycopersici*, and *F. solani* (teleomorph *Nectria haematococca*) were on max score were analyzed by SMART.

### A series of *FvLCP1* mutants tested for FB_1_ biosynthesis

A *FvLCP1* gene knock-out mutant (ΔFvlcp1) was generated to test FB_1_ production (F1g. S1 A). Homologous recombination was confirmed by PCR (data not shown) and Southern analysis (Fig. S1 B). When compared with the wild-type progenitor, ΔFvlcp1 was indistinguishable when grown on PDA and V8 agar media (Fig. S1 C). This outcome suggested that this gene is not critical for *F. verticillioides* vegetative growth. To further study the physiological roles of *FvLCP1*, particularly its key predicted functional motifs, on FB_1_ production, we generated ΔFvlcp1 complementation strain as well as FvLcp1 motif-deletion mutants ΔSP, ΔLysM and ΔChtBD1, under its native promoter and terminator (Fig 4A). HPLC analysis were performed to elucidate the production of the FB_1_ on maize cracked kernels by these strains. Strains ΔFvlcp1 and ΔLysM showed most drastic reduction in FB_1_ compared to the wild-type strain, *Δ*SP, and ΔChtBD1 mutants (Fig. 4B). These results suggested that *FvLCP1* gene is important for FB_1_ production. Gene complementation with full *FvLCP1* construct restored FB_1_ production to the wild-type level. Specifically, the LysM domains was determined to be important for wild-type level FB_1_ production in *F. verticillioides*.

**Figure 4.**
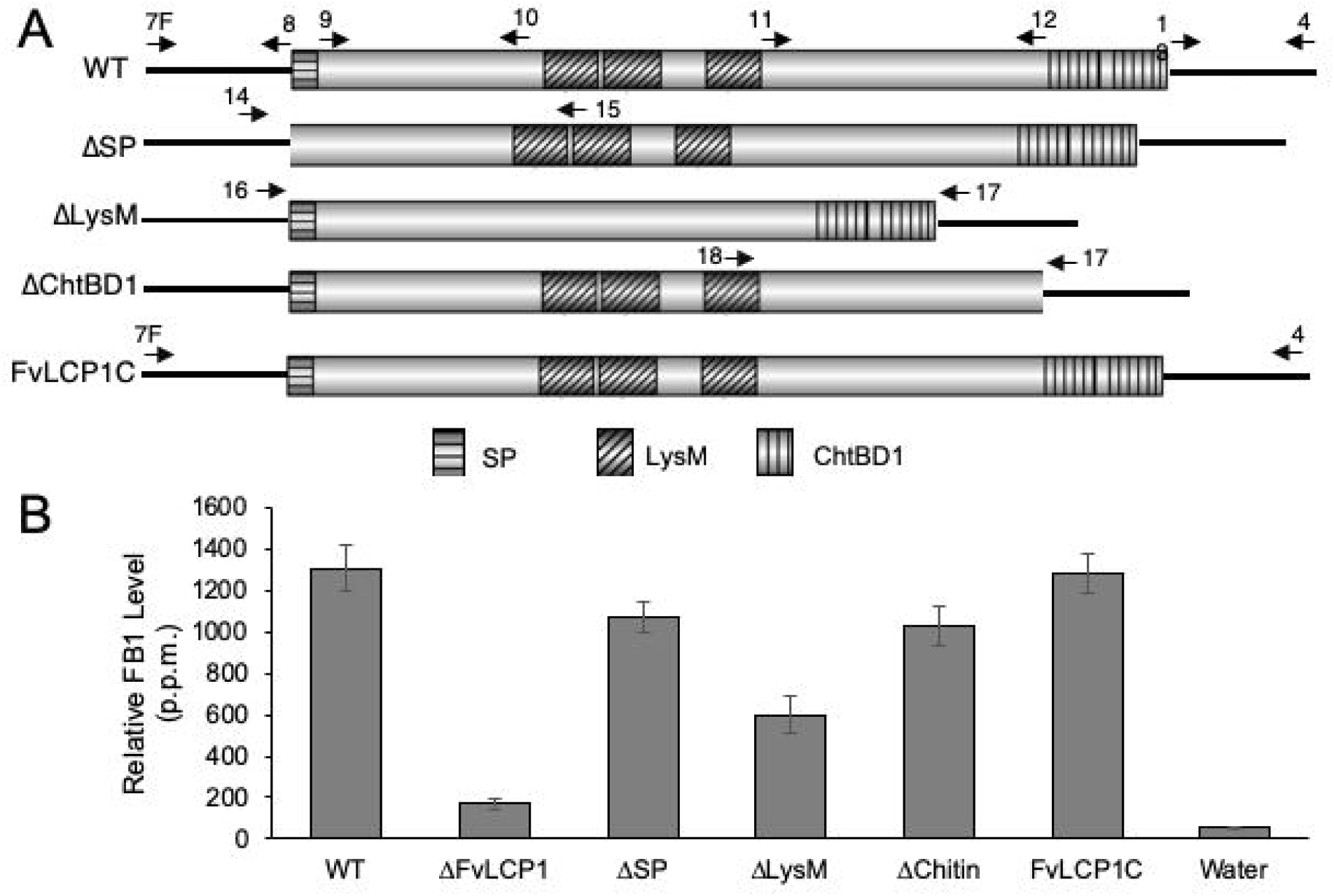
Schematic overview of *FvLCP1* locus manipulation showed LysM motif is important for fumonisin production. (A) Four complementation constructs were generated via homologous recombination; ΔSP) complete deletion of signal peptide deletion, (ΔLysM) complete deletion of LysM domains, (ΔChtBD1) complete deletion of ChtBD1 domains, and FvLCP1C) complete wild-type gene. These constructs were all driven by its native promoter and terminator. Complementation constructs used to complement ΔFvlcp1 gene deletion strain. Arrows above the WT schematic model represent the primers used to amplify the specific DNA fragments to generated all the above constructs. Arrows above the other schematic models represent primers used for PCR screening analysis. (B) Quantification of fumonisin B1 (FB1) production in *F. verticillioides* strains. 2×10^6^ spores of WT, ΔFvLCP1, ΔSP, ΔLysM, Δ ChtBD1 and FvLCP1C strains were inoculated on nonviable autoclaved maize kernels and incubated for 7 days at 25 °C under a 14-h light/10-h dark cycle. FB_1_ production was quantified by high-performance liquid chromatography (HPLC) analysis. FB_1_ biosynthesis was normalized to growth with ergosterol contents. All values represent the means of three biological replications with standard errors shown as error bars.

### FvLcp1 is a secreted protein in *F. verticillioides*

Recent studies have demonstrated that proteins targeted for secretion, notably effectors and effector-like proteins in fungi, usually has signal peptide in the N-terminal region. Signal peptide is composed of a short, positively charged amino-terminal region (n-region), a central hydrophobic region (h-region), and a more polar carboxy-terminal region (c-region), which is targeted and cleaved by the signal peptidase (Von Heijine., 1990; Von Heijine., 1998). Our *in silico* analyses using SignalP 4.1 (Petersen et al., 2011) showed that FvLcp1 has one signal peptide and one cleavage site. Prediction of subcellular localization by ProtComp v6.0 software (http://www.softberry.com/) suggested that this protein is an extracellular secreted protein (Fig. S2 A).

To further experimentally verify that FvLcp1 is a secreted protein, we performed yeast secretion trap assay. The full-length open reading frames of FVEG_01584 and one well known secreted protein *Magnaporthe oryzae* AvrPiz-t with or without signal peptides were cloned into pYST-0. Yeast transformants expressing the *FvLcp1^1-2202^*:*SUC2* or the *avrPiz-t ^1-327^*:*SUC2* fusions grew on sucrose, whereas yeast expressing the *FvLcp1^31-2202^*:*SUC2* or the *avrPiz-t^25-327^*:*SUC2* fusions, the same proteins without the predicted SPs, failed to grow (Fig. S2 B). The results demonstrated that FvLcp1 is a protein secreted extracellularly in *F. verticillioides*.

### The LysM domains in FvLcp1 group together with fungal specific LysM clade rather than previously identified fungal effectors

In plant pathogenic fungi, recent studies have shown that proteins with LysM motifs (including Ecp6, Mg3LysM, Slp1, ChElp1, ChElp2 and Vd2LysM) can bind chitin and suppress chitin-induced plant innate immune response (de Jonge *et al*., 2010; Marshall *et al*., 2011; Mentlak *et al*., 2012; Takahara *et al*., 2016; Kombrink *et al*., 2017). To test the phylogenetic relationship between FvLcp1 and other characterized LysM containing proteins in fungi and plants, we performed a comparative LysM sequence analysis. Consistent with a previous report (Kombrink *et al*., 2017), all LysM sequences were divided into two distinct clades (Fig. 5). The first clade was formed by ‘fungal-specific’ LysMs, which might be involved in fungal development rather than pathogen-host interactions (Seidl-Seiboth *et al*., 2013; Kombrink *et al*., 2017). The second clade was formed by LysMs from fungal effector and LysMs from plant receptors (de Jonge *et al*., 2010; Marshall *et al*., 2011; Liu, B *et al*., 2012; Mentlak *et al*., 2012; Takahara *et al*., 2016; Kombrink *et al*., 2017). When compared with sequences from ‘effector/receptor’ LysMs, the ‘fungal-specific’ LysMs show distinct conserved sequences with three highly conserved cysteine residues. Based on available information, we concluded that LysM domains found in FvLcp1 belong to the first ‘fungal-specific’ clade.

**Figure 5.**
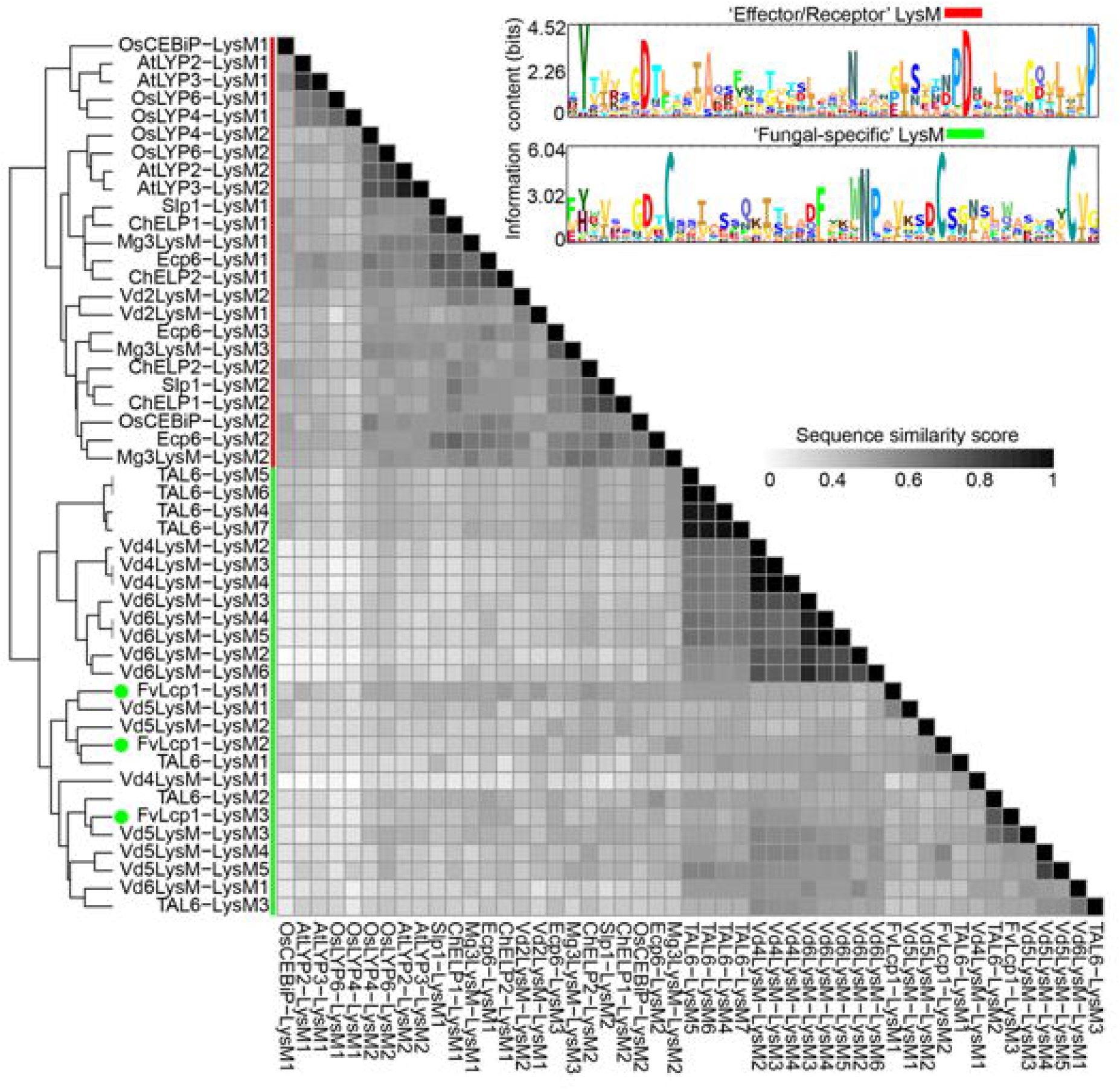
Comparative analysis of LysM domains in fungi and plants. Sequence similarities between different LysMs from fungi and plants are visualized by the heat map. Two clades were performed by hierarchically clustered. Red and green lines indicated the two separate clades. Green dots represent the presence of three different LysM domains, respectively. Numbers after LysM indicates the order of different LysM domains from N-terminus to C-terminus in disparate proteins. In the upper right corner, the two sets of conserved sequences are displayed by the sequence logo. A Red rectangle indicated the sequence logo of Effector/Receptor’ LysM domains. The fungal-specific’ LysM sequence logo was emphasized by a green rectangle.

### Subcellular localization of FvLcp1

*FvLCP1* with its native promoter was inserted into pKNTG vector, which contains a GFP tag and the geneticin-resistance marker. We verified the accuracy of the construct by PCR and sequencing, before transforming the construct into *F. verticillioides* wild-type strain. One geneticin-resistance transformant, which expressed FvLcp1 and fused C-terminal GFP under the native promoter, was selected for further observation. We also stained *F. verticillioides* cells with FM4-64 (vacuolar membrane dye) or CMAC (vacuolar dye) (Fig. 6A). We were able to observe FvLcp1 signals near plasma membrane, septum and vacuole in conidia, germinated conidia, and vegetative hyphae stages. Co-localization analyses confirmed that CMAC epifluorescence was observed along with FvLcp1-GFP signal on vacuole. FvLcp1-GFP was also surrounded by FM4-64 epifluorescence which outlined the vascular membrane (Fig. 6B).

**Figure 6.**
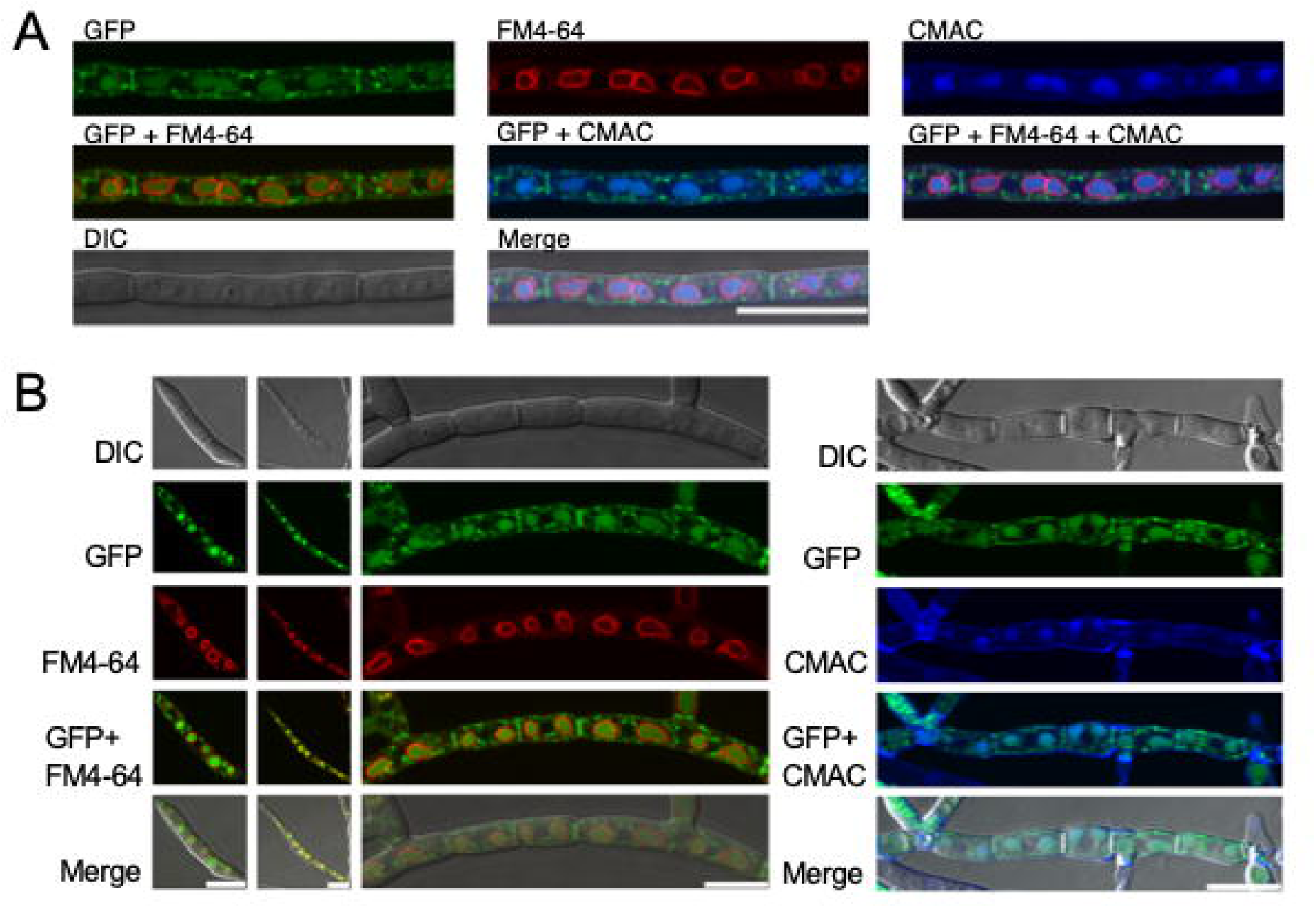
FvLcp1mainly localizes to plasma membrane, septum and vacuole in the vegetative growth stage. A) Subcellular localization of FvLcp1-GFP in vegetative hyphae. The cells were stained with FM4-64 and CMAC. Bar=10 μm. B) Subcellular localization of FvLcp1-GFP in conidia, germinated conidia, and vegetative hyphae. The cells were stained with FM4-64 and CMAC. Bars=10 μm.

### FvLcp1 is accumulated in appressorium structure in the early infection stage of *M. oryzae*

Maize kernel and *F. verticillioides* is not the most accessible host-pathogen system to conduct experiment to study fungal protein movement and localization. Therefore we adopted rice -*M. oryzae*, a well-advanced model system for studying host-pathogen interactions, for our study (Ebbole, 2007). By infecting rice sheath cells, we can visualize the action of interested proteins in the early plant infection process (Mosquera *et al*., 2009). To determine the localization and role of FvLcp1 in the early infection stage of the pathogen, we transferred FvLcp1-GFP into the rice blast pathogen *M. oryzae.* We showed that Guy11/FvLcp1-GFP mainly localized in appressorium structure (Fig. 7A), which is an infectious structure and generates enormous turgor to penetrate the plant cuticle (Howard *et al*., 1991). However, we learned that FvLcp1 does not further enter rice as *M. oryzae* develop appressoria and invasive hyphae. This result suggest that FvLcp1 may play a role in host recognition and assist in penetration but not in host colonization in *M. oryzae*. When we compared virulence in Guy11 and Guy11/FvLcp1-GFP transformants by performing spray inoculation assays on susceptible rice cultivar CO39, we did not observe a significant difference between wild-type Guy11 and Guy11 expressing FvLcp1-GFP (Fig. 7B), suggesting that the addition of FvLcp1 in *M. oryzae* did not enhance fungal virulence.

**Figure 7.**
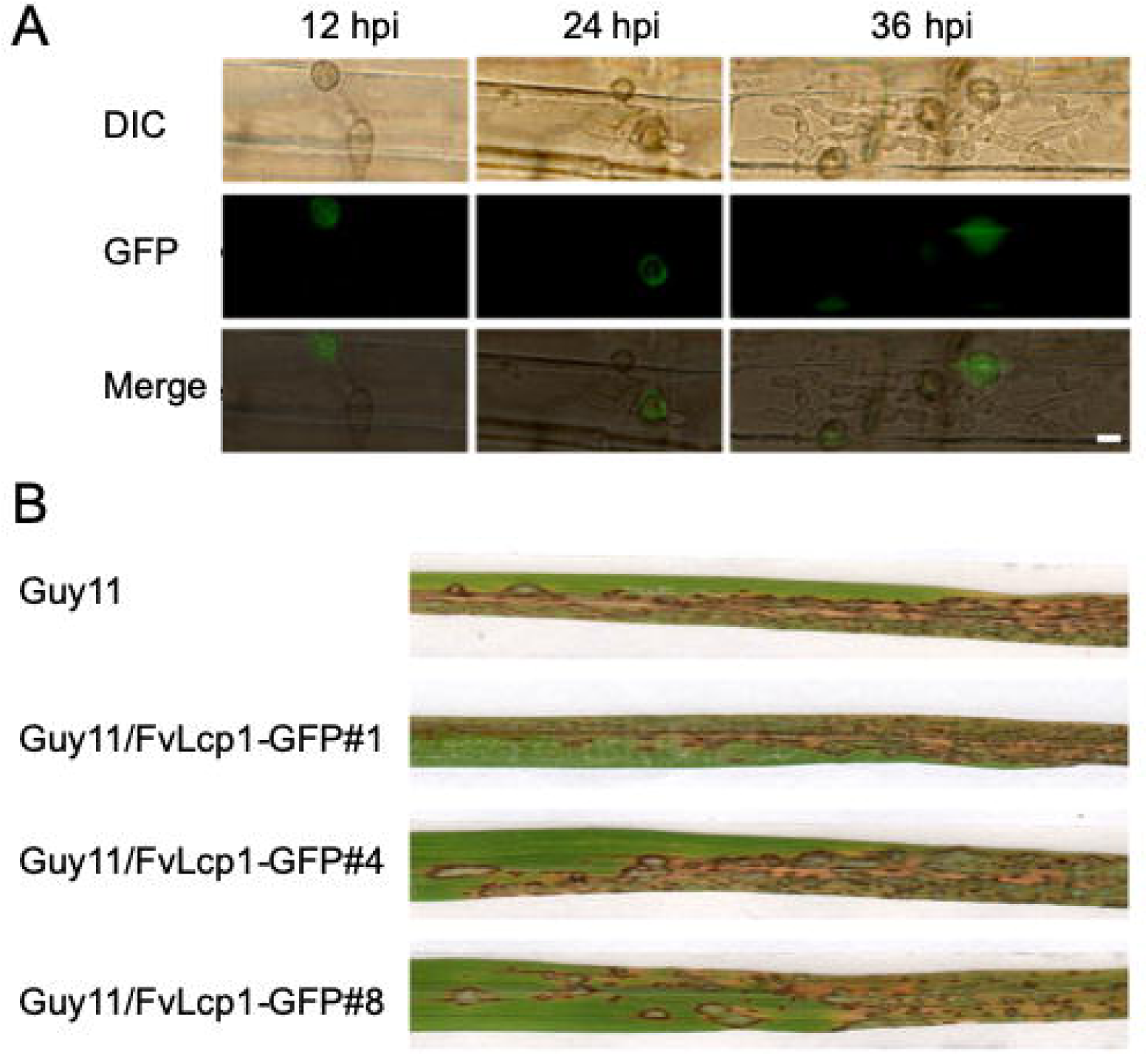
FvLcp1 localizes in appressorium structure in the early infection stage of *M. oryzae.* (A) Subcellular Localization of FvLcp1-GFP in *M. oryzae* Guy11 (Guy11/FvLcp1-GFP) during biotrophic growth on susceptible rice sheath epidermal cells at 12hpi, 24 hpi and 36hpi. Green fluorescence signals were detected in appressorium structure. Bar=10 μm. (B) Guy11/ FvLcp1-GFP shows similar blast disease symptom with *M. oryzae* wild-type Guy11. Susceptible rice cultivar CO39 were sprayed with conidial suspensions with equal concentration (1×10^5^ spore/mL in 0.02% Tween 20 solution).

### FvLcp1 is able to bind chitin via ChtBD1 domains

FvLcp1 contains LysM and ChtBD1 domains which have been reported to bind chitin (de Jonge *et al*., 2010; Marshall *et al*., 2011; Liu, Z *et al*., 2012; Mentlak *et al*., 2012; Liu *et al*., 2016; Takahara *et al*., 2016; Kombrink *et al*., 2017), and therefore we experimentally tested whether FvLcp1 is a chitin binding protein. First, N-terminal Glutathione S-transferase (GST)-tagged mature FvLcp1 (without its signal peptide) protein was heterologously produced in *E. coli* strain Rosetta. Chitin magnetic beads were incubated with mature FvLcp1, and we observed mature FvLcp1 in the insoluble pellet fractions which indicates mature FvLcp1 as a chitin binding protein that can coprecipitate with insoluble chitin magnetic beads. To investigate the roles of LysM and ChtBD1 in FvLcp1 chitin binding, we generated GST-tagged FvLcp1ΔLysM and FvLcp1ΔChtBD1 proteins (Fig. 8A). When compared with the mature FvLcp1 protein, FvLcp1ΔChtBD1 could not be found in the pellet fraction which shows FvLcp1ΔChtBD1 did not precipitate with chitin magnetic beads. However, FvLcp1ΔLysM was detected in the pellet fractions, exhibiting strong affinity as mature FvLcp1. Overall, our data indicated that FvLcp1 is able to bind chitin, and ChtBD1 domains are essential for this activity.

**Figure 8.**
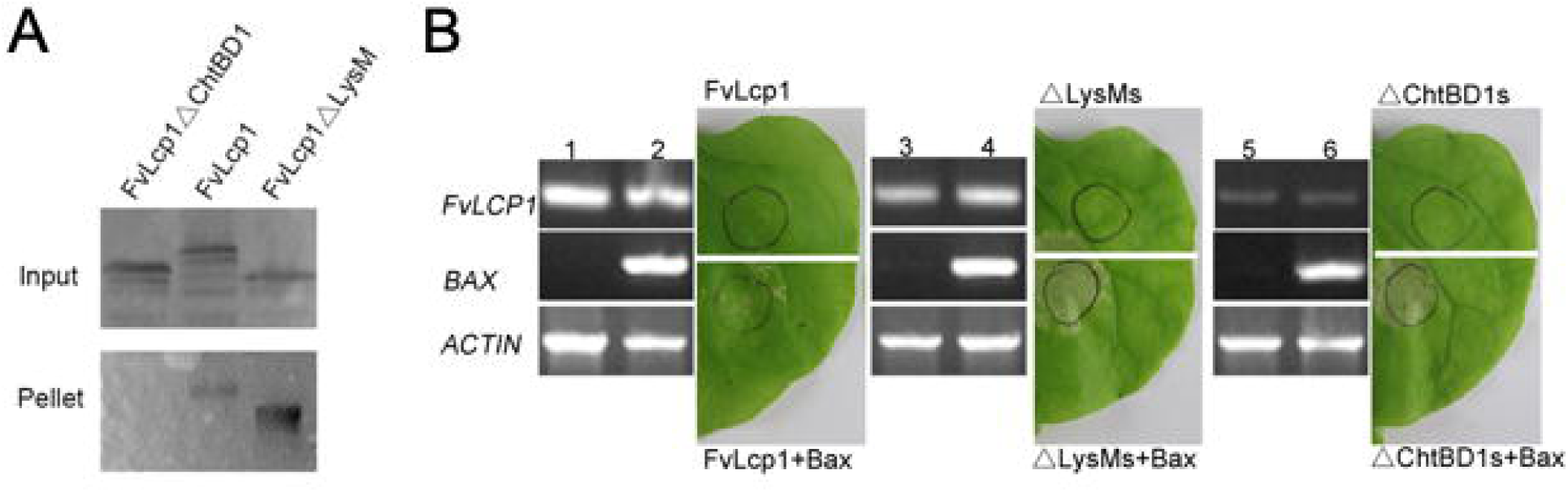
FvLcp1 is able to bind chitin via ChtBD1 domains suppresses the BAX triggered hypersensitive plant cell death. (A) Mature FvLcp1 protein and truncated FvLcp1 proteins (ΔLysMs and ΔChtBD1s) produced in *E. coli* were tested for their chitin binding ability by affinity precipitation with the chitin magnetic beads. The input (total proteins) and pellet fraction were both analyzed. Detection of mature FvLcp1 and FvLcp1ΔLysMs in the pellet indicates FvLcp1 binds to chitin by ChtBD1 domains rather than LysM domains. (B) Symptoms of transient of Mature FvLcp1 and truncated FvLcp1 in *N. benthamiana* leaves by agroinfiltration approach. Top panel images show the representative infectious results for mature FvLcp1 protein and two truncated FvLcp1 proteins. The middle panel models indicate the order of infiltrations in the leaves of *N. benthamiana*. The bottom panel RT-PCR results confirmed the expression of mature FvLcp1, two truncated FvLcp1 proteins, and BAX. RT-PCR results from 1) leaves infiltrated with mature FvLcp1; 2) leaves infiltrated with mature FvLcp1 and BAX; 3) leaves infiltrated with mature FvLcp1 deleting LysM domains; 4) leaves infiltrated with mature FvLcp1 deleting LysM domains and BAX; 5) leaves infiltrated with mature FvLcp1 deleting ChtBD1 domains; 6) leaves infiltrated with mature FvLcp1 deleting ChtBD1 domains and BAX.

### FvLcp1 suppresses the BAX triggered hypersensitive plant cell death but requires both LysM and ChtBD1 domains

Many plant pathogen effectors suppress and interfere with plant innate immune response and trigger susceptibility (Jones & Dangl, 2006; Li *et al*., 2009; Dong *et al*., 2015; Li *et al*., 2015; Sharpee *et al*., 2017). For example, AvrPiz-t, an avirulence protein from *M. oryzae* can suppress BAX-induced plant cell death in *N. benthaminan* (Li *et al*., 2009). While our earlier in silico study revealed that FvLcp1 is a fungal-specific LysM protein, we were interested in testing whether mature FvLcp1 (without signal peptide) could interfere with the plant cell death triggered by BAX. We infiltrated *N. benthaminan* leaves with *A. tumefaciens* carrying mature FvLcp1. We chose AvrPiz-t and empty vector (pGWB505) as positive and negative controls, respectively (Li *et al*., 2009). As previously reported, AvrPiz-t can suppress the BAX-triggered plant cell death. Our results showed that mature FvLcp1 alone could not induce plant cell death. However, FvLcp1 was able to suppress plant cell death triggered by BAX (Fig. 8B). We further tested LysM and ChtBD1 domain-specific deletion mutants to further demonstrate the roles of LysM domains and ChtBD1 domains in the suppressing BAX. Transient expression experiments showed that LysM deletion mutant and ChtBD1 deletion mutant both failed to suppress the plant cell death induced by BAX (Fig. 9B). In order to investigate whether the mature FvLcp1 protein interferes with BAX expression, we performed RT-PCR to detect the expression of BAX and FvLcp1. RT-PCR confirmed that FvLCP1 and BAX were both expressed in *N. benthamiana* leaves. These results indicated that FvLcp1 is able to suppress the BAX-triggered plant cell death, but both LysM and ChtBD1 domains are required to suppress BAX-induced plant cell death.

## DISCUSSION

In this study, our aim was to identify potential subnetwork modules differentially activated in *F. verticillioides* when inoculated on two different maize inbred B73 and hybrid 33k44 kernels. Our premise was that maize pathogen *F. verticillioides* utilizes unique mechanisms for host organelle adaptation, virulence, and mycotoxin biosynthesis and that there will be subtle differences when challenging kernel with different genetic background. To predict reliable *F. verticillioides* functional subnetwork modules and to capture dynamic changes in gene association during kernel colonization and mycotoxin production, we proposed a more robust systematic network-based comparative analysis approach using RNA-seq data. Specifically, we were focused on genes that encode putative secreted proteins that may play a role in *F. verticillioides* – kernel association. We have modified our previous analytical strategy (Kim *et al*., 2018b) by first selecting differentially expressed genes that encode proteins harboring signal peptide. Through this approach, we identified four potential *F. verticillioides* virulence-associated subnetwork modules where the member genes were harmoniously coordinated and significantly differentially activated between two maize kernels. We selected one subnetwork module with the predicted hub FVEG_01584 for further characterization, particularly focusing on whether protein encoded by this hub gene can be secreted into plant host and subsequently impact kernel rot disease and fumonisin biosynthesis. We were intrigued by the fact that FVEG_01584 (designated *FvLCP1*) encoded a putative secreted protein with three LysM domains and two chitin-binding (ChtBD1) domains.

The LysM motif (Pfam PF01476) was first described in bacteriophage lysome which is responsible for cleaving the bacterial cell wall (Garvey *et al*., 1986). Besides lysomes, LysMs can also be found in chitinases, receptor-like kinases, glycoside hydrolases, transglycosylases, peptidases, amidases in both prokaryotic and eukaryotic proteins (Ponting *et al*., 1999; Buist *et al*., 2008). The LysM motif consists of around 50 amino acids, in which the first 16 residues and final 10 residues are conserved, and more than 400 putative fungal LysM domain-containing proteins have been identified (de Jonge & Thomma, 2009). In fungal pathogen *Cladosporium fulvum*, LysM effector Ecp6 can bind to chitin directly and compete with chitin receptor from the plant host, thereby repressing the chitin-induced resistance (de Jonge *et al*., 2010). Another LysM effector Slp1, which is the ortholog of Ecp6 in rice blast pathogen *M. oryzae*, is up-regulated during the early infection stage of *M. oryzae* (Mentlak *et al*., 2012). In order to promote infection, Slp1 competes with chitin elicitor binding protein (CEBiP) in rice (*Oryza sativa*) by directly binding to chitin oligosaccharides released from fungal cell wall (Mentlak *et al*., 2012). Moreover, chitin binding domain is considered to be involved in binding or recognition of chitin sub units (Butler *et al*., 1991). In wheat pathogen *Mycosphaerella graminicola,* LysM proteins Mg1LysM and Mg3LysM were shown to play important role in protecting fungal hyphae against chitinases secreted by competitors. In addition, Mg3LysM mutant strains were severely impaired in leaf colonization, lesion formation, and asexual sporulation (Marshall *et al*., 2011; Lee *et al*., 2014). Recently, *Colletotrichum higginsianum* extracellular LysM proteins ChElp1 and ChElp2 were found to play dual roles in appressorial function and suppression of chitin-triggered plant immunity (Takahara *et al*., 2016). Notably, LysM effector Vd2LysM in *Verticillium dahliae*, a soil-borne wilt pathogen, was also reported to suppresses chitin-induced immune responses and protects hyphae against degradation by plant hydrolytic enzymes (Kombrink *et al*., 2017).

All these identified LysM proteins do not contain any recognizable motif other than LysM. These proteins were classified into type A LysM proteins which constitutes over 70% of fungal LysM proteins (de Jonge & Thomma, 2009). The other LysM proteins were found to contain LysM domains in combination with other enzymatic domains, such as chitin binding domain (found in types B and D), the CVNH domain (found in type C), and the amidase domain (found in type E). These domains have been proposed to have an important role in carbohydrate binding. Thus far, the LysM proteins harboring these additional enzymatic domains were not extensively studied. The only protein of this classification that was reported is CVNH-LysM, a type-C LysM protein, in rice blast fungus *M. oryzae*. A study showed that CVNH and LysM domains, while structurally intact and functionally competent, function independently for carbohydrate binding and play crucial role in host infection (Koharudin *et al*., 2011). In *F. verticillioides,* we identified FvLcp1 that contains a signal peptide and three LysM domains in the N-terminus and two chitin-binding domains in the C-terminus, indicating that this is a type D fungal LysM protein. To date, no functional study has been published on type D fungal LysM protein. One of the questions we asked in this study was whether LysM domains and chitin binding domains carry out distinct roles when compared with entire FvLcp1 protein. Our result demonstrated that the LysM domains play an important role in fumonisin production, while chitin-binding domains are important for binding chitin. Significantly, our study also showed that full FvLcp1 protein is necessary for suppression of the BAX-triggered hypersensitive plant cell death when tested in *N. benthamiana*.

As discussed earlier, LysM proteins also bind chitin. Chitin is a major component of fungal cell walls and plays an important role in fungal - host interactions (Kombrink *et al*., 2011; Rovenich *et al*., 2016). Chitin-triggered immunity is known to cause induction of pathogenesis-related genes, and we therefore sought to examine the effect of FvLcp1 on induction of maize defense gene expression (Fig. S3). WT and FvLcp1 mutant conidia were inoculated on surface-sterilized live maize kernels. Expression of key maize defense genes (*ZmLOX5, ZmLOX10, OPR8, PR1*) was tested 6 and 8 days-post-inoculation. *F. verticillioides* WT induced *ZmLOX5* expression 3.5-folds and 1.6-folds higher than *Fvlcp1* mutant on 6 days-post-inoculation and 8 days-post-inoculation samples, respectively. However, there was no significant difference between the treatments for *OPR8* gene expression, and *LOX10* and *PR1* expression studies were inconclusive. *ZmLOX5* is a lipoxygenase which plays important roles in plant defense against pathogens (De La Fuente *et al*., 2013), and this result showed that FvLcp1 is involved in triggering *ZmLOX5* mediated defense pathway. Clearly, further experiments are necessary to better understand how FvLcp1 impacts defense response in hosts. There is also a need to identify a putative receptor of FvLcp1 in maize and the signaling pathway which will help to explain how FvLcp1 triggers the plant immune response.

One of the intriguing questions we asked while conducting this study was whether FvLcp1 that can be secreted into plant cells and trigger specific host responses in maize kernels. Since experimenting this idea was not easily achievable using maize kernels, we performed our study in *M. oryzae*-rice and *A. tumefaciens-N. benthamiana* systems. *M. oryzae* is one of the most intensively studied plant pathogens, particularly when we discuss molecular host-pathogen associations and effector biology (Ebbole, 2007; Yoshida *et al*., 2009; Mentlak *et al*., 2012; Liu *et al*., 2013; Dong *et al*., 2015). Most of the identified effectors in *M. oryzae* encode small secreted proteins (<200 amino acids) and do not have any known protein domains (Li *et al*., 2009; Wu *et al*., 2015). However, not all fungal effectors share characteristics found in *M. oryzae* effectors. As we discussed earlier, LysM effectors such as Ecp6, which bind chitin by LysM domains (de Jonge *et al*., 2010). *FvLCP1* encodes a large secreted protein of 733 amino acids and contains three LysM domains and two ChtBD1 domains. However, our chitin binding assay confirmed that two ChtBD1 domains in FvLCP1 are responsible for chitin binding rather than the three LysM domains. Transient expression testing in *N. benthamiana* with BAX showed that FvLcp1 could suppress the BAX-induced plant cell death, suggesting that FvLcp1 is involved in suppressing plant defense response. When we expressed FvLcp1 in *M. oryzae*, the protein was localized to appressoria initially suggesting that FvLcp1 could be involved in host recognition and penetration. However, spray assay showed no clear difference between Guy11 and Guy11/FvLcp1-GFP, demonstrating that the presence of FvLcp1 adds no further benefit in virulence in the rice blast pathogen. In similar fashion, the deletion mutants of BAS effectors (BAS1, BAS2 and BAS3) in *M. oryzae* did not result in loss-of-pathogenicity phenotypes (Mosquera *et al*., 2009). In necrotrophic fungi, plant cell death could promote infection and colonization process (Wang *et al*., 2014). However, our study showed FvLcp1, as a secreted *F. verticilioides* protein, could serve as a suppressor of plant cell death rather than an inducer of cell death, which potentially suggests that suppressing host cell death could play a critical benefit to its infection process. Based on the role of FvLcp1 in FB_1_ production, it can be hypothesized that the fungus utilizes FvLcp1 to sustain the viability of plant host in the early infection stage. This is the first report where a secreted protein, particularly a type-D LysM protein with chitin-binding domain, in mycotoxigenic fungus *F. verticillioides* is potentially involved in suppressing host cell death to perhaps maximize its ability to produce fumonisin while it initiates kernel infection. There are a series of intriguing studies that are required to better understand *F. verticillioides*-maize interactions, particularly on relationship between plant cell death and FB_1_ production.

## ACKNOWLEDGEMENTS

This research was supported in part by the Agriculture and Food Research Initiative Competitive Grants Program (2013-68004-20359) from the USDA National Institute of Food and Agriculture and by National Science Foundation China (Project No.31601599). We would also like to thank Chinese Scholarship Council (201500090070) for Mr. Jun Huang. We would like to thank Dr. Yijuan Han, Ms. Xiaojie Zhang, Ms. Mingyue Shi, and Mr. Lianyu Lin (Fujian Agricultural and Forestry University, China) for their assistance during the data collection. Authors declare no conflict of interest.

## AUTHOR CONTRIBUTIONS

H.Z., M.K., J.H. and W.B.S. conceived and designed the experiments. H.Z., M.K., J.H., H.Y., T.Y., L.S. and W.Y. performed the experiments. H.Z, M.K, J.H., W.Y. and W.B.S. analyzed the data. H.Z., M.K., J.H., and W.B.S. wrote the paper.

