## Supplementary Methods for "FvLcp1, a type-D fungal LysM protein with Chitin-binding domains, is a secreted protein involved in host recognition and fumonisin production in *Fusarium verticillioides* - maize kernel interaction"

### Supplementary Method

#### A. RNA-Seq Preprocessing

We preprocessed the RNA-Seq datasets by following steps; i) alignment: RNA-Seq reads were mapped to the *F. verticillioides* reference genome provided by the BROAD Institute (http://www.broadinstitute.org) using TopHat2 (Kim *et al.*, 2013) and subsequently quantified to be counts for all genes by HTSeq (Anders *et al.*, 2015); ii) filtering for relatively insignificantly expressed genes was completed as described in our previous study (Kim *et al.*, 2018a); iii) Three different datasets (**PGEM: Preprocessed Gene Expression Matrix**) with three different combinations of two-step normalization; i) **PGEM-1**: RPKM (Mortazavi *et al.*, 2008) - control; ii) **PGEM-2**: RPKM (Mortazavi *et al.*, 2008) -TMM (Robinson *et al.*, 2010); iii) **PGEM-3**: TPM (Wagner *et al.*, 2012)-TMM (Robinson *et al.*, 2010); were obtained by normalizing the gene expression levels in within-replicates and across-replicates, respectively. For normalization using control genes, the mean expression levels of beta-tubulin genes in *F. verticillioides* (FVEG_04081 and FVEG_05512) were applied.

#### B. Seed genes

Since we targeted potential effectors, we collected genes that contained signal peptide through UniProt (www.uniprot.org) among the preprocessed genes, thereby narrowing down all *F. verticillioides* genes to 1235 genes (7.78%). Among the collected genes, we selected not only top 10 significantly differentially expressed *F. verticillioides* genes in maize B73, but also top 10 significantly differentially expressed *F. verticillioides* genes in maize 33K44 based on *t*-test statistics scores as well as F scores (generated by one-way ANOVA) of the 1235 genes. The selected genes showed relatively high *t*-test statistics score*s* in all three PGEMs and also relatively significant F scores in at least one of the PGEMs.

**C. Co-expression networks**

We inferred *F. verticillioides* co-expression networks on the two maize strains (B73 and 33K44) for all three PGEMs to establish fundamental association between genes as applied in our previous studies (Kim *et al.*, 2015; Kim *et al.*, 2018a; Kim *et al.*, 2018b). Edges (*i.e.*, connections) were generated between genes when two genes held significant strength of association (*i.e.*, at least certain partial correlation coefficients (Hero & Rajaratnam, 2012)), where we constructed five different co-expression networks with five distinct partial correlation thresholds to weaken dependency on a specific threshold cut-off level for each PGEM. Sizes of the networks were classified based on numbers of edges as they gradually enlarged from the smallest (*i.e.*, 400,000 edges) to the largest (*i.e.*, 2,000,000 edges).

**D. Initial subnetwork module extension**

For each PGEM, the initial module extension was performed as described in our previous study (Kim *et al.*, 2018a). Based on a *F. verticillioides* co-expression network, we began screening for candidate subnetwork modules having genes up to two. From a seed gene, we extended edges to every neighboring gene and evaluated every discriminative power of probabilistic subnetwork activity levels of the modules. As explained in our previous work [3], we predicted activity level $\boldsymbol{\delta}(\boldsymbol{e})$ of a subnetwork module by supposing $\boldsymbol{\zeta} = \{\boldsymbol{g_1}, \boldsymbol{g_2}, \boldsymbol{g_3}, \dots, \boldsymbol{g_n}\}$, a group of genes in a subnetwork module, and $\boldsymbol{e} = \{\boldsymbol{e^1}, \boldsymbol{e^2}, \boldsymbol{e^3}, \dots, \boldsymbol{e^n}\}$, expression levels of the given genes. The subnetwork module activity level $\boldsymbol{\delta}(\boldsymbol{e})$ was estimated as follows

$$\boldsymbol{\delta(e)} = \sum_{\boldsymbol{k=1}}^{\boldsymbol{n}} \boldsymbol{log}\ [\frac{\boldsymbol{f_1^k(e^k)}}{\boldsymbol{f_2^k(e^k)}}]$$

where $\boldsymbol{log}\ [\frac{\boldsymbol{f_1^k(e^k)}}{\boldsymbol{f_2^k(e^k)}}]$ is the log-likelihood ratio (LLR), and $\boldsymbol{f_1^k(e)}$ and $\boldsymbol{f_2^k(e)}$ are the conditional probability density function (PDF) in the two maize kernels (B73 vs 33K44). Then, the discriminative power of the given module between the two conditions was computed based on *t*-test statistics. Next, we allowed a module with the optimal discriminative power enhance of module activity level to be a candidate module, and also two additional suboptimal modules with two specific conditions: i) minimum discriminative power enhance of probabilistic activity level of an extended module should exceed at least 5%, ii) discriminative power difference of subnetwork activity levels between the optimum and the sub-optimum should be less than 2%.

**E. Subnetwork module with adaptive branch-out**

While the initial step followed our previous one, the current proposed network-based comparative analysis approach is an updated methodology. After the initial screening for subnetwork modules up to two member genes, we started our adaptive branch-out technique and optimized subnetwork modules so they can be differentiated as much as possible between the two strains (maize B73 vs maize 33K44). In this adaptive branching-out, we permitted relatively lower discriminative power enhance in early stage by considering relatively small number of possible new members of early stage as well as relatively high *t*-test scores of the seed genes. Also, we adaptively searched for new member genes by adjusting stopping criterion (**MDPE: Minimum Discriminative Power Enhance**) by considering following requirements for candidate subnetwork modules; i) relatively high discriminative power of a given module between the two strains; ii) relatively large number of connections between seed genes and other member genes; iii) significant GO term annotation of member genes in a given module; iv) maximum distance of four between the seed gene and all member genes in a given module. The significance of GO terms was considered by *p*-value of the Benjamini-Hochberg false discovery rate (FDR) approach (Benjamini & Hochberg, 1995) provided by g:Profiler (http://biit.cs.ut.ee/gprofiler/) (Reimand *et al.*, 2011). The GO terms were taken into account when at least 30% of the member genes were associated with one term and the *p*-values were lower than 0.05. The update of stopping criterion (MDPE) $\boldsymbol{t}$ at each extension is as follows

$$\boldsymbol{t_{m+1} = (1+\beta)t_m}$$

where

$$\boldsymbol{\beta = 0.05 \quad (m = 1)}$$

$$\boldsymbol{\beta = \gamma * \frac{\sum_{m=1}^{m-1} \ln t_m}{\sum_{m=1}^{m} \ln t_m} \quad (m \geq 2)}$$

where $m$ is the number of member genes of a subnetwork module and $\beta$ is adaptive factor of stopping criterion update and also $\gamma$ is the desired (maximum) stopping criterion, initially applied as 0.01 (meaning 10%). Since this adaptive branching-out computes the next stopping criterion based on the previous stopping criteria (MDPE values), it relatively further increases the next stopping criterion if the previously updated stopping criteria are relatively low and vice versa.

##### F. Identification of subnetwork module

After collecting all candidate subnetwork modules through our adaptive branch-out approach for all three PGEMs, we applied our post-pruning process to identify relatively more robust functional subnetwork modules by considering edge activity level. In order to estimate it, we took into account three criteria; i) how significantly it differentiated its module on one strain from the other strain; ii) how easily and repeatedly it demonstrated strong association with other genes including its seed gene across different co-expression networks based on different PGEMs; iii) whether it held connections between member genes for functional coherence (*i.e.*, GO terms). We first selected one representative subnetwork module with the highest discriminative power from each PGEM and combined the three representative modules into a module. In this merging process, we created a score matrix, $\boldsymbol{A}$, where $\boldsymbol{a_{ij}}$ is an element of $\boldsymbol{A}$ based on subnetwork module member genes, $\boldsymbol{\zeta} = \{\boldsymbol{g_1}, \boldsymbol{g_2}, \dots, \boldsymbol{g_s}, \dots, \boldsymbol{g_n}\}$, and we gave a point to $\boldsymbol{a_{ij}}$ when $\boldsymbol{g_i}$ and $\boldsymbol{g_j}$ had a connection to each other as follows.

$$\boldsymbol{A}(\boldsymbol{s}) = \begin{cases} \mathbf{1}, & \boldsymbol{if} \ \sum_{n} \boldsymbol{a_{ij}^{n}} \ (\boldsymbol{i} \neq \boldsymbol{j}) \ \geq \ \mathbf{2} \\ \mathbf{0}, & \boldsymbol{if} \ \sum_{n} \boldsymbol{a_{ij}^{n}} \ (\boldsymbol{i} \neq \boldsymbol{j}) \ = \ \mathbf{1} \end{cases}$$

where $\boldsymbol{s} = \boldsymbol{i}\ \boldsymbol{or}\ \boldsymbol{j}$ when $\boldsymbol{g_s}$ is considered. In order to measure impact of each gene (*i.e.*, $\boldsymbol{g_s}$) on its module differentiation between the two strains, we computed discriminative power difference $\boldsymbol{D}(\boldsymbol{s})$ between both activity levels (one with the gene and the other without the gene) as follows.

$$\boldsymbol{T}(\boldsymbol{s}) = \begin{cases} \mathbf{1}, & \boldsymbol{if} \ \boldsymbol{D}(\boldsymbol{s}) = [\boldsymbol{\delta}(\boldsymbol{e})]_{\zeta}^{t-test\ score} - [\boldsymbol{\delta}(\boldsymbol{e})]_{\zeta-\{g_s\}}^{t-test\ score} \ > \ \mathbf{0} \\ \mathbf{0}, & \boldsymbol{if} \ \boldsymbol{D}(\boldsymbol{s}) = [\boldsymbol{\delta}(\boldsymbol{e})]_{\zeta}^{t-test\ score} - [\boldsymbol{\delta}(\boldsymbol{e})]_{\zeta-\{g_s\}}^{t-test\ score} \ < \ \mathbf{0} \end{cases}$$

We took into consideration potential functional relevance of a gene (*i.e.*, $\boldsymbol{g_s}$) by considering GO terms as follows.

$$\boldsymbol{G}(\boldsymbol{s}) = \begin{cases} \mathbf{1}, & \boldsymbol{if} \ \boldsymbol{g_s} \ \boldsymbol{is\ annotated\ to\ significant\ GO\ term\ with\ other\ member\ genes} \\ \mathbf{0}, & \boldsymbol{otherwise} \end{cases}$$

where *p*-value of GO term significance ≤ 0.05. Based on the three criteria, we finally evaluated edge activity level of a gene (*i.e.*, $\boldsymbol{g_s}$) defined as follows.

$$\boldsymbol{EA}(\boldsymbol{s}) = \boldsymbol{A}(\boldsymbol{s}) + \boldsymbol{max}[\boldsymbol{T}(\boldsymbol{s}) \cdot \boldsymbol{G}(\boldsymbol{s})]$$

After computing all edge activity levels ($\boldsymbol{EA}$) in the combined module, we started eliminating edges whose $\boldsymbol{EA}$ was 0 from outside. When a gene with $\boldsymbol{EA} = \mathbf{0}$ from outer side was removed, edge activity levels of all module member genes were computed again. The post-pruning process continuously executed until all module member genes contained edges whose $\boldsymbol{EA}$ was larger than 0.

#### G. *Nucleic acid manipulation, polymerase chain reaction (PCR), and transformation*

Briefly, for ΔSP construct generation, primers 1F and 2R were used for SP 5' flanking region generation, and primers 3F and 4R were used for SP 3' flanking region generation. 5' and 3' flanking region fragments were then fused by single-joint PCR by using primers 1F and 4R (Fig. 4A). Similarly, ΔLysM and ΔChtBD1 constructs were generated. FvLCP1C construct was amplified by primers 7F and 4R. Then we introduced each of the constructs together with a geneticin-resistance (*GEN*) marker into the Δ*Fvlcp1* mutant protoplasts. All primers used in this study were listed in Supplementary Method H. *F. verticillioides* protoplast were generated and transformed following standard protocol (Sagaram & Shim, 2007).

#### H. Primers used in this stud

| Primer Name | 5'-3' sequence |
|---|---|
| 1F | TCC CGA CAG ACA GAC GAA TG |
| 2R | TAG ATG CCG ACC GGG AAC ACC GAA AAT GCC GTA TGT CTT |
| 3F | CCA CTA GCT CCA GCC AAG AGC TTT GGC TCG GGA TGT AC |
| 4R | AGT TAG CAG GAA TCG TGG TGG |
| 5F | AGAAACAGCCTACGCAGACCT |
| 6R | CTGTTGGCCAGAGGCACATA |
| 7F | TGC AGA AGC TAC CAA GTC GCAG |
| HYG-F | CTTGGCTGGAGCTAGTGGAGGTCAA |
| HY-R | GTATTGACCGATTCCTTGCGGTCCGAA |
| YG-F | GATGTAGGAGGGCGTGGATATGTCCT |
| HYG-R | GTTCCCGGTCGGCATCTACTCTAT |
| 8 | 01584-SP-R: GACTAACGCCGACAATAACCA |

| | |
|---|---|
| 9 | 01584-SP-F: TTATTGTCGGCGTTAGTCGATATGGAGCTTGATACCGCTGA |
| 10 | 01584-LysM-R: CGAACTGGCGGAAGTGTAGA |
| 11 | 01584-LysM-F: TACACTTCCGCCAGTTCGAGCATTTCCCTCGATGACTTC |
| 12 | 01584-Chitin-R: AAGGGTCTCGTGGCAGTAGAA |
| 13 | 01584-Chitin-F: TACTGCCACGAGACCCTTTGTTGCTCGCAGTACGGTTAC |
| 14 | SP-SCREEN-F: AGGTAAATCAGTCTTTCGACC |
| 15 | SP-SCREEN-R: TTGTGTACTCGGAACTACAGC |
| 16 | LysM-screen-F: AAGACATACGGCATTTTCGGT |
| 17 | LysM-screen-R: AGTACATCCCGAGCCAAAGCT |
| 18 | Chitin-screen-F: AGCATTTCCCTCGATGACTTC |
| PL1F | AGGAACCCAATCTTCAAAAAGACATACGGCATTTTCGGT |
| PL1R | GTACATCCCGAGCCAAAGCT |
| PL2F | CCTCGTTTCTGCTGTTATGTC |
| PL2R | CAAAGGGCTGTCTGGATGT |
| LYSM_GFPF | CCGCTCGAGTTTCAGCCATTTGGGAAATTA |
| LYSM_GFPR | CCCATCGATAGTACACACTCCATACTGCTTTTG |
| PL3F | CTGGTTATTGTCGGCGTTAGTC |
| PL3R | GGTCTGCGTAGGCTGTTTCTTAT |
| PL4F | GGGGACAAGTTTGTACAAAAAAGCAGGCTATGGAGGATCTCTTTTCACTTGGTGAA |
| PL4R | GGGGACCACTTTGTACAAGAAAGCTGGGTCCTAAGTACACACTCCATACTGCTT |
| PL5F | GGGGACAAGTTTGTACAAAAAAGCAGGCTATGGAGGATCTCTTTTCACTTGGTGAA |
| PL5R | GGGGACCACTTTGTACAAGAAAGCTGGGTCAGTACACACTCCATACTGCTTTTG |
| 01584-LysM-F | TACACTTCCGCCAGTTCGAGCATTTCCCTCGATGACTTC |
| 01584-LysM-R | CGAACTGGCGGAAGTGTAGA |
| 01584-Chitin-F | TACTGCCACGAGACCCTTTGTTGCTCGCAGTACGGTTAC |
| 01584-Chitin-R | AAG GGT CTC GTG GCA GTA GAA |
| fvlcp1-GST-F | TTCTAGACTCCATGGGTCGACTTGAGGATCTCTTTTCACTTGGTGAAAT |
| fvlcp1-GST-R | TCAGTCACGATGAATAAGCTTCTAAGTACACACTCCATACTGCTTTTGG |
| BAX-RT2F | TTTTGCTACAGGGTTTCATCCA |
| BAX-RT2R | CCCGAAGTAGGAGAGGAGGC |
| NbACT-F | ATGGCAGACGGTGAGGATATTCA |
| NbACT-R | GCCTTTGCAATCCACATCTGTTG |
| LCP1 YZ4F | ATGGACCTGAGGAGGACGAATG |
| LCP1 YZ4R | CCCGAGGAGACGGAGGAACT |

Underlined sequences represent overlaps used for joint PCR.
