## Supplementary Tables for "FvLcp1, a type-D fungal LysM protein with Chitin-binding domains, is a secreted protein involved in host recognition and fumonisin production in *Fusarium verticillioides* - maize kernel interaction"

**Table S1**. General statistics of RNA-Seq datasets

| | Maize 33K44 | | Maize B73 | |
|---|---|---|---|---|
| | 6 dpi | 8 dpi | 6 dpi | 8 dpi |
| Type of run | paired-end | | | |
| Read length | 125 (bp) | | | |
| Mean # of reads aligned | 7155626 | 13473461 | 6532043 | 18301912 |
| Median depth of coverage | 505.3 | 951.4 | 461.3 | 1292.4 |

**Table S2**. Co-expression network size

A. Statistics of the three co-expression networks on maize B73

| # of edge | PGEM 1 (RPKM-beta) | | PGEM 2 (RPKM-TMM) | | PGEM 3 (TPM-TMM) | |
|---|---|---|---|---|---|---|
| | Threshold | # of genes | Threshold | # of genes | Threshold | # of genes |
| Size-1 ≈ 400,000 | 0.7620 | 13524 | 0.7320 | 13522 | 0.7300 | 13522 |
| Size-2 ≈ 800,000 | 0.7170 | 13532 | 0.6860 | 13532 | 0.6845 | 13533 |
| Size-3 ≈ 1,200,000 | 0.6860 | 13532 | 0.6550 | 13533 | 0.6525 | 13533 |
| Size-4 ≈ 1,600,000 | 0.6610 | 13532 | 0.6300 | 13533 | 0.6280 | 13533 |
| Size-5 ≈ 2,000,000 | 0.6400 | 13532 | 0.6090 | 13533 | 0.6075 | 13533 |

B. Statistics of the three co-expression networks on maize 33K44

| # of edge | PGEM 1 (RPKM-beta) | | PGEM 2 (RPKM-TMM) | | PGEM 3 (TPM-TMM) | |
|---|---|---|---|---|---|---|
| | Threshold | # of genes | Threshold | # of genes | Threshold | # of genes |
| Size-1 ≈ 400,000 | 0.7375 | 13524 | 0.7270 | 13522 | 0.7290 | 13522 |
| Size-2 ≈ 800,000 | 0.6915 | 13532 | 0.7010 | 13532 | 0.7025 | 13533 |
| Size-3 ≈ 1,200,000 | 0.6640 | 13532 | 0.6740 | 13533 | 0.6755 | 13533 |
| Size-4 ≈ 1,600,000 | 0.6355 | 13532 | 0.6450 | 13533 | 0.6465 | 13533 |
| Size-5 ≈ 2,000,000 | 0.6140 | 13532 | 0.6043 | 13533 | 0.6055 | 13533 |

**Table S3**. Percentage of common edges on co-expression networks between PGEMs

| # of edge | PGEM-1 <-> PGEM-2 | PGEM-1 <-> PGEM-3 | PGEM-2 <-> PGEM-3 |
|---|---|---|---|
| Size-1 ≈ 400,000 | 64.5% | 62.7% | 94.6% |
| Size-2 ≈ 800,000 | 67.8% | 65.8% | 94.8% |
| Size-3 ≈ 1,200,000 | 69.7% | 68.1% | 95.0% |
| Size-4 ≈ 1,600,000 | 71.2% | 69.4% | 95.2% |
| Size-5 ≈ 2,000,000 | 72.4% | 70.5% | 95.3% |

**Table S4**. *t*-test statistics score and F score (ANOVA) of the selected seed genes

| *F. verticillioides* genes with signal peptide significantly differentially expressed in B73 | | | | | | |
|---|---|---|---|---|---|---|
| Gene | *t*-test score | | | F score | | |
| | PGEM-1 | PGEM-2 | PGEM-3 | PGEM-1 | PGEM-2 | PGEM-3 |
| FVEG_01584 | 7.4 | 4.1 | 4.1 | 4.5 | 1.2 | 1.1 |
| FVEG_01729 | 4.9 | 5.7 | 5.7 | 3.6 | 6.3 | 6.9 |
| FVEG_01761 | 6.5 | 5.3 | 5.4 | 24.1 | 3.4 | 3.6 |
| FVEG_07670 | 7.1 | 5.2 | 5.2 | 24.6 | 24.8 | 26.6 |
| FVEG_12539 | 6.9 | 5.7 | 5.6 | 2.3 | 27.6 | 28.8 |
| FVEG_14015 | 7.2 | 5.6 | 5.6 | 21.4 | 29.7 | 27.3 |
| FVEG_04110 | 4.0 | 4.5 | 4.1 | 4.0 | 0.1 | 0.2 |
| FVEG_05442 | 4.6 | 4.8 | 4.8 | 16.0 | 2.4 | 2.1 |
| FVEG_08308 | 4.0 | 4.0 | 4.2 | 10.4 | 4.9 | 4.9 |
| FVEG_11722 | 5.7 | 4.2 | 4.3 | 12.1 | 0.4 | 0.5 |

| *F. verticillioides* genes with signal peptide significantly differentially expressed in 33K44 | | | | | | |
|---|---|---|---|---|---|---|
| Gene | *t*-test score | | | F score | | |
| | PGEM-1 | PGEM-2 | PGEM-3 | PGEM-1 | PGEM-2 | PGEM-3 |
| FVEG_05692 | 10.3 | 7.9 | 7.7 | 6.7 | 4.9 | 5.1 |
| FVEG_09285 | 7.6 | 7.7 | 7.7 | 12.7 | 2.9 | 2.5 |
| FVEG_09653 | 7.0 | 8.2 | 8.3 | 6.2 | 1.3 | 1.2 |
| FVEG_11787 | 10.3 | 8.8 | 8.6 | 2.7 | 5.7 | 6.1 |
| FVEG_13513 | 9.3 | 10.2 | 9.9 | 0.3 | 16.7 | 20.4 |
| FVEG_13534 | 9.9 | 14.4 | 14.9 | 5.0 | 1.7 | 1.5 |
| FVEG_00397 | 7.2 | 7.5 | 7.4 | 9.6 | 11.4 | 10.4 |
| FVEG_01726 | 6.3 | 7.9 | 7.8 | 5.2 | 0.3 | 0.1 |
| FVEG_05574 | 7.1 | 6.8 | 6.7 | 4.9 | 8.2 | 8.8 |
| FVEG_09387 | 8.1 | 7.3 | 7.2 | 14.2 | 6.2 | 6.0 |

**Table S5**. Number of subnetwork modules generated based on the selected seed genes

| B73 | | | | 33K44 | | | |
|---|---|---|---|---|---|---|---|
| Gene | PGEM-1 | PGEM-2 | PGEM-3 | Gene | PGEM-1 | PGEM-2 | PGEM-3 |
| FVEG_01584 | 1 | 4 | 2 | FVEG_05692 | 0 | 1 | 0 |
| FVEG_01729 | 0 | 0 | 0 | FVEG_09285 | 0 | 4 | 2 |
| FVEG_01761 | 0 | 1 | 2 | FVEG_09653 | 0 | 0 | 1 |
| FVEG_07670 | 1 | 2 | 1 | FVEG_11787 | 0 | 0 | 0 |
| FVEG_12539 | 0 | 0 | 0 | FVEG_13513 | 0 | 0 | 0 |
| FVEG_14015 | 0 | 0 | 0 | FVEG_13534 | 0 | 1 | 1 |
| FVEG_04110 | 0 | 0 | 0 | FVEG_00397 | 2 | 5 | 3 |
| FVEG_05442 | 0 | 0 | 0 | FVEG_01726 | 1 | 2 | 3 |
| FVEG_08308 | 0 | 0 | 0 | FVEG_05574 | 2 | 5 | 2 |
| FVEG_11722 | 2 | 2 | 4 | FVEG_09387 | 0 | 0 | 0 |
