## Supplementary Figures for "FvLcp1, a type-D fungal LysM protein with Chitin-binding domains, is a secreted protein involved in host recognition and fumonisin production in *Fusarium verticillioides* - maize kernel interaction"

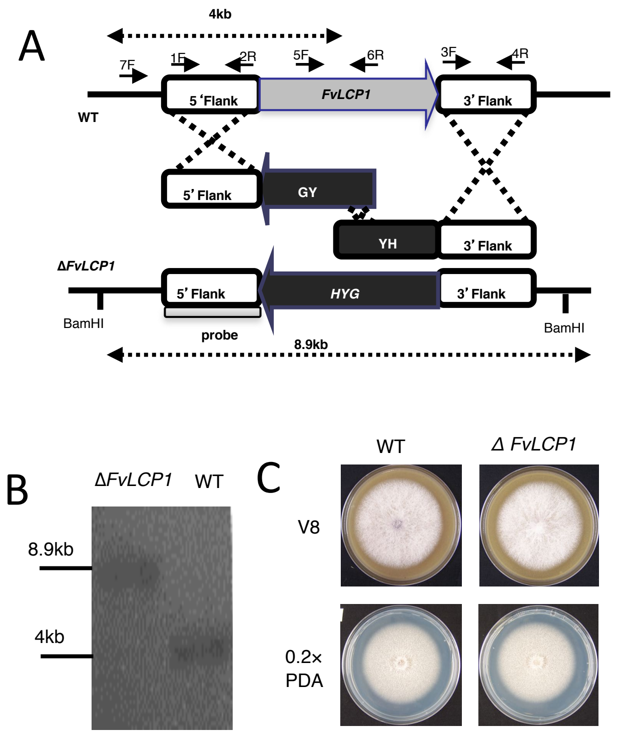


**Figure S1. Schematic representation of the *FvLCP1* disruption complementation strategies and growth phenotype on solid agar plate.** (A) Target replacement of *FvLCP1* with the hygromycin phosphotransferase gene *(HPH)* by the split marker technique through homologous recombination Arrows indicate primers used for PCR. *HYG*, hygromycin phosphotransferase gene, *HY*, *HYG* 5' partial amplicon, *YG*, *HYG* 3' partial amplification. (B) Southern analyses of wild-type (WT) and knockout mutant *(ΔFvLCP1)*. 5'-flanking region was used as a probe for Southern hybridization. (C) Vegetative growth of WT and *ΔFvLCP1 w*ere examined on V8 and 0.2XPDA agar plates. Strains were point inoculated with an agar block (0.5 cm in diameter) and incubated for 6 days at 25 °C under 14 h light/10 h dark cycle.

Zhang et al. Fig. S2

A

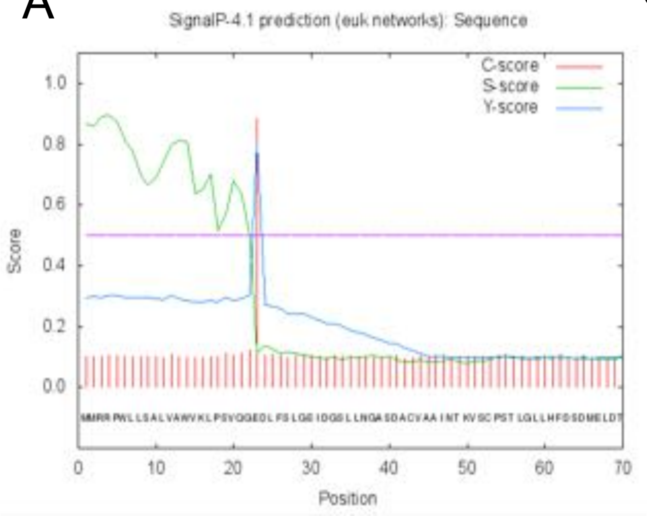


C

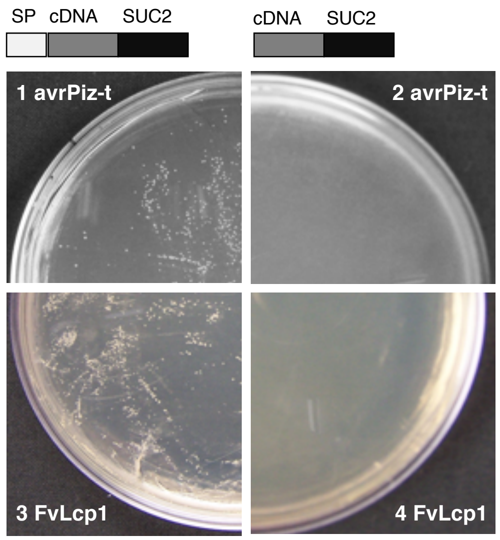


B

```
5712 multiple located sequences are accepted
ProtComp Version 9.0. Identifying sub-cellular location (Animals&Fungi)
Seq name:  test sequence, Length=733
Significant similarity by DBSCAN-P - NONE
Predicted by Neural Nets - Extracellular (Secreted) with score    2.6
Integral Prediction of protein location: Extracellular (Secreted) with score    8.9
Location weights:     LocDB / PotLocDB / Neural Nets / Pentamers / Integral
 Nuclear               0.0 /      0.0 /        0.04 /       0.00 /     0.12
 Plasma membrane       0.0 /      0.0 /        0.24 /       0.37 /     0.74
 Extracellular         0.0 /      0.0 /        2.63 /       1.88 /     8.90
 Cytoplasmic           0.0 /      0.0 /        0.00 /       0.00 /     0.13
 Mitochondrial         0.0 /      0.0 /        0.00 /       0.23 /     0.00
 Endoplasm. retic.     0.0 /      0.0 /        0.00 /       0.00 /     0.00
 Peroxisomal           0.0 /      0.0 /        0.00 /       0.00 /     0.00
 Lysosomal             0.0 /      0.0 /        0.00 /       0.02 /     0.00
 Golgi                 0.0 /      0.0 /        0.09 /       0.80 /     0.11
 Vacuolar              0.0 /      0.0 /        0.00 /       0.00 /     0.00
```

**Figure S2. FvLcp1 is a secreted protein in *F. verticillioides*.** (A) *In silico* analyses using SignalP 4.1 showed that FvLcp1 has one signal peptide and one cleavage site. (B) Prediction of subcellular localization by ProtComp v6.0 software (http://www.softberry.com/) suggested that this protein is an extracellular secreted protein. (C) Yeast secretion trap assay to experimentally confirm physical secretion. YST analysis of *Magnaporthe oryzae* avrPiz-t and *F. verticillioides* FvLcp1 with (panels 1 and 3, respectively) or without (panels 2 and 4, respectively) their SPs. Yeast secretion trap (pYST0) vectors to identify FvLcp1 as a secreted protein. YST analysis of *Magnaporthe oryzae* avrPiz-t and *F. verticillioides* FvLcp1 with (panels 1 and 3, respectively) or without (panels 2 and 4, respectively) their SPs.

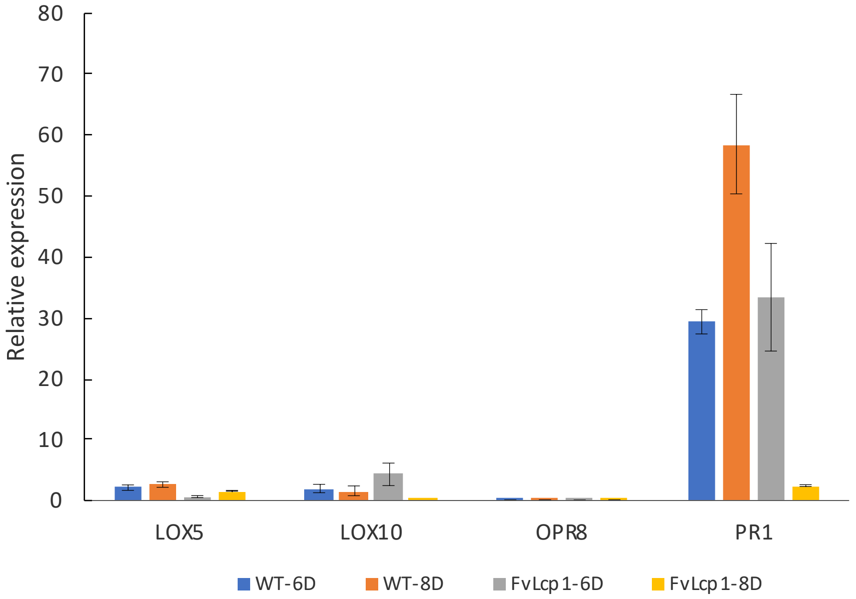


**Fig. S3. Examination of the effect of FvLCP1 gene deletion on induction of maize defense-related genes.** WT and FvLcp1 mutant conidia were inoculated on surface-sterilized live maize kernels. Relative expression of key maize defense genes (*ZmLOX5, ZmLOX10, OPR8, PR1*) was tested 6 and 8 days-post-inoculation. The maize GAPDH gene was used as the reference.
